# A Phylogenomic Framework, Evolutionary Timeline, and Genomic Resources for Comparative Studies of Decapod Crustaceans

**DOI:** 10.1101/466540

**Authors:** Joanna M. Wolfe, Jesse W. Breinholt, Keith A. Crandall, Alan R. Lemmon, Emily Moriarty Lemmon, Laura E. Timm, Mark E. Siddall, Heather D. Bracken-Grissom

**Affiliations:** Division of Invertebrate Zoology & Sackler Institute of Comparative Genomics, American Museum of Natural History, New York, NY 10024, USA; Department of Earth, Atmospheric & Planetary Sciences, Massachusetts Institute of Technology, Cambridge, MA 02139, USA; Museum of Comparative Zoology & Department of Organismic & Evolutionary Biology, Harvard University, Cambridge, MA 02138, USA; Florida Museum of Natural History, University of Florida, Gainesville, FL 32611, USA; RAPiD Genomics, Gainesville, FL 32601, USA; Computational Biology Institute, The George Washington University, Ashburn, VA 20147, USA; Department of Invertebrate Zoology, National Museum of Natural History, Smithsonian Institution, Washington, DC 20012, USA; Department of Scientific Computing, Florida State University, Dirac Science Library, Tallahassee, FL 32306, USA; Department of Biological Science, Florida State University, Tallahassee, FL 32306, USA; Department of Biological Sciences, Florida International University, North Miami, FL 33181, USA

**Keywords:** Decapoda, Pancrustacea, Crustacea, Phylogenomics, Anchored Hybrid Enrichment, Systematics

## Abstract

Comprising over 15,000 living species, decapods (crabs, shrimp, and lobsters) are the most instantly recognizable crustaceans, representing a considerable global food source. Although decapod systematics have received much study, limitations of morphological and Sanger sequence data have yet to produce a consensus for higher-level relationships. Here we introduce a new anchored hybrid enrichment kit for decapod phylogenetics designed from genomic and transcriptomic sequences that we used to capture new high-throughput sequence data from 94 species, including 58 of 179 extant decapod families, and 11 of 12 major lineages. The enrichment kit yields 410 loci (>86,000 bp) conserved across all lineages of Decapoda, eight times more molecular data than any prior study. Phylogenomic analyses recover a robust decapod tree of life strongly supporting the monophyly of all infraorders, and monophyly of each of the reptant, ‘lobster’, and ‘crab’ groups, with some results supporting pleocyemate monophyly. We show that crown decapods diverged in the Late Ordovician and most crown lineages diverged in the Triassic-Jurassic, highlighting a cryptic Paleozoic history, and post-extinction diversification. New insights into decapod relationships provide a phylogenomic window into morphology and behavior, and a basis to rapidly and cheaply expand sampling in this economically and ecologically significant invertebrate clade.

## 1. Introduction

Decapod crustaceans, broadly categorized into ‘shrimp’, ‘lobsters’, and ‘crabs’, are embedded in the public consciousness due to their importance as a global food source worth over $24 billion [1]. Several ornamental species are also popular in the pet trade [2,3], and some lobsters and crayfish may be promising models for cancer and aging research [4]. Furthermore, decapods are a major faunal component of a bewildering variety of global habitats, including the open ocean, seafloor vents and seeps, caves, coral reefs, mangroves and estuaries, intertidal mud and sand, freshwater streams and lakes, semi-terrestrial locations, and in symbiosis with other animals (**Figure 1**). Decapods have diversified over the course of 455 million years resulting in over 15,000 living and 3,000 fossil species recognized in approximately 233 families [5,6]. Despite the economic and ecological significance of the clade, higher-level phylogenetic relationships among decapods have proven recalcitrant.

**Figure 1.**
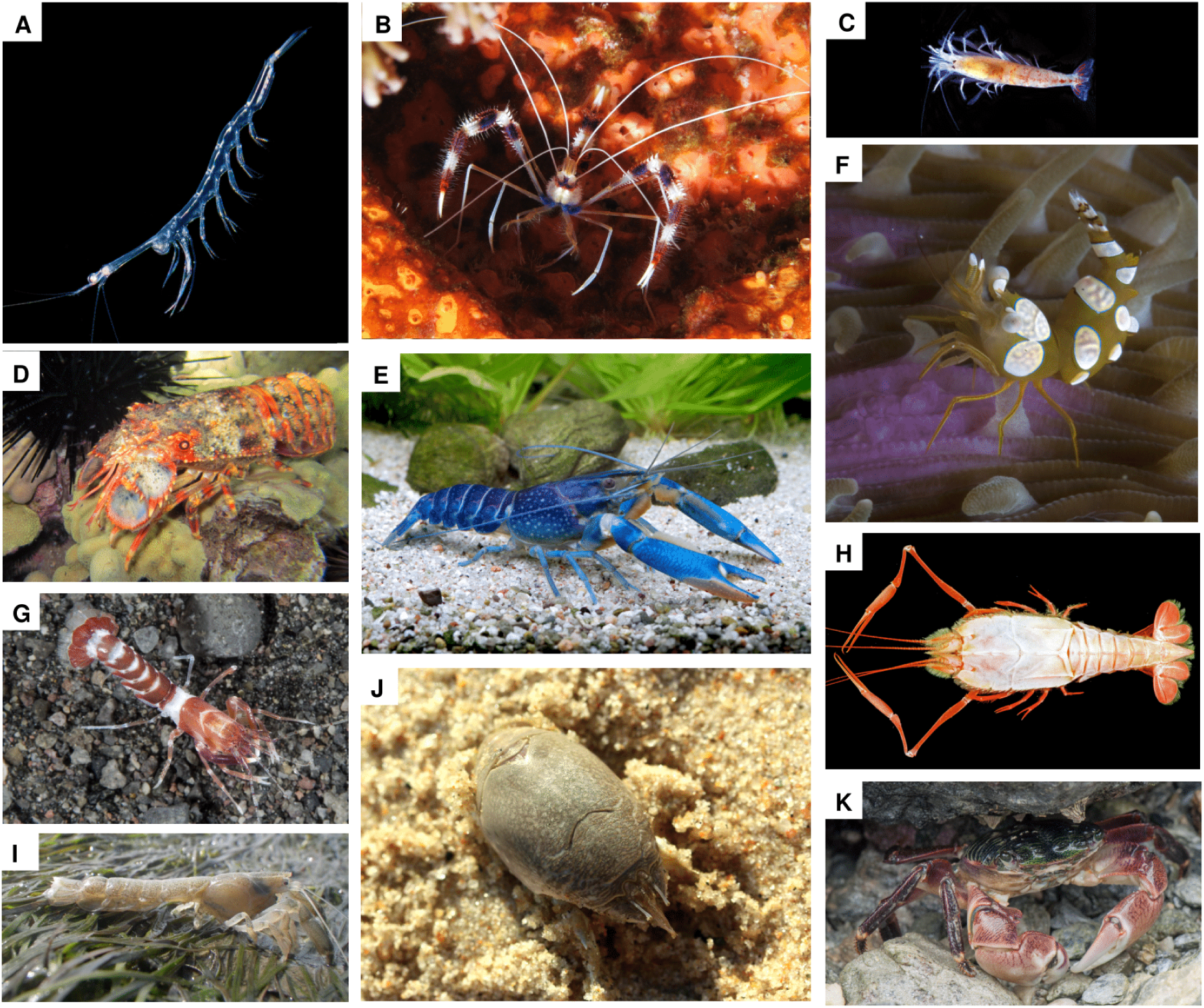
Representatives of major decapod lineages. *(a) Lucifer* sp. (Southeast Florida, USA) (Dendrobranchiata); *(b) Stenopus hispidus* (Komodo, Indonesia) (Stenopodidea); *(c) Procaris chacei* (Bermuda) (Procarididea); *(d) Arctides regalis* (Maui, Hawaii, USA) (Achelata); *(e) Cherax quadricarinatus* (aquarium specimen) (Astacidea); *(f) Thor amboinensis* complex (Ternate, Maluku Islands, Indonesia) (Caridea); *(g) Axiopsis serratifrons* (Bali, Indonesia) (Axiidea); *(h) Stereomastis sculpta* (specimen ULLZ 8022) (Polychelida); *(i) Upogebia* cf. *pusilla* (Arcachon Bay, France) (Gebiidea); *(j) Emerita talpoida* (Westerly, Rhode Island, USA) (Anomura); *(k) Pachygrapsus crassipes* (Catalina Island, California, USA) (Brachyura). Photo credits: *(a)* L. Ianniello; *(b)* A. Vasenin, license CC-BY-SA; *(c)* T.M. Iliffe; *(d*, *k)* J. Scioli; *(e)* C. Lukhaup; *(f)* C.H.J.M. Fransen; *(g)* A. Ryanskiy; *(h)* D.L. Felder; *(i)* X. de Montaudouin; *(j)* J.M. Wolfe.

The majority of work is restricted to studies using morphology [7–9], up to nine targeted mitochondrial and nuclear genes [6,10–19], and more recently complete mitogenomes of 13 genes [20–24]. Mitogenomic data can be problematic for reconstructing ancient nodes [25], and indeed, deeper relationships receive poor support [24]. As part of a larger analysis, decapods were included in a recent transcriptomic study [26], but with limited taxon sampling within the order. This plurality of results, several based on the same underlying data [25], have reported conflicting deep relationships among decapods. Without a robust phylogeny, comparative inferences about morphology, development, ecology, and behavior are limited.

Herein, phylogenomic sequencing of nuclear genes is leveraged for the first time in decapods, using anchored hybrid enrichment (AHE), a technique previously applied to vertebrates [27], plants [28], and clades of terrestrial arthropods that have diverged at least 100 Myr more recently than decapods [29–32]. Anchored Hybrid Enrichment specifically targets conserved coding regions that are flanked by less conserved sequence regions, with the goal of optimizing phylogenetic informativeness at multiple levels of divergence [27]. Unlike popular transcriptomic approaches, AHE does not require fresh or specially preserved tissues (critical for sampling the diversity of decapods, since many lineages are rare, confined to the deep sea, and/or have complicated life histories). Instead, AHE allows the use of ethanol-preserved specimens; however prior genomic and/or transcriptomic data are required to determine genomic target regions.

Here we combine new genomic and transcriptomic sequences to build AHE probes spanning all of Decapoda, ultimately sequencing 86 species and 7 outgroups. The enrichment kit we constructed can easily be used by the systematics community for future studies of decapod evolution. Ours is the first example of a strongly supported phylogenomic analysis including almost all major decapod lineages sequenced for over 400 loci, the largest dataset yet compiled for this group. With the inclusion of 19 vetted fossil calibration points, we also present the first divergence time analysis incorporating a well-supported topology for the entire decapod clade.

## 2. Methods

### (a) Probe design

Target AHE loci were identified using our previous workflows (e.g. [29,31,32]; **Figure S1**) at the FSU Center for Anchored Phylogenomics (www.anchoredphylogeny.com). Targets were based on genomic resources from 23 decapod species (**Table S1**), including nine newly sequenced genomes (~6-31x coverage; **Table S2**) and four newly sequenced transcriptomes (**Table S3-S4**). Details of genome and transcriptome sequencing in **Extended Methods 1a-c**. Best-matching reads were identified in the two highest-recovery taxa (**Table 1**, RefsA), as well as reference sequences from the red flour beetle *Tribolium castanaeum* [29,32], resulting in 823 preliminary AHE target sequences. As in Hamilton et al. [29], we screened exemplar transcriptomes from five major decapod lineages (**Table 1**, RefsB) for the best-matching transcript, then aligned in MAFFT v7.023 [33], requiring representation in at least four of the lineages and resulting in 352 final targets. Additional details in **Extended Methods 1d**.

**Table 1.**
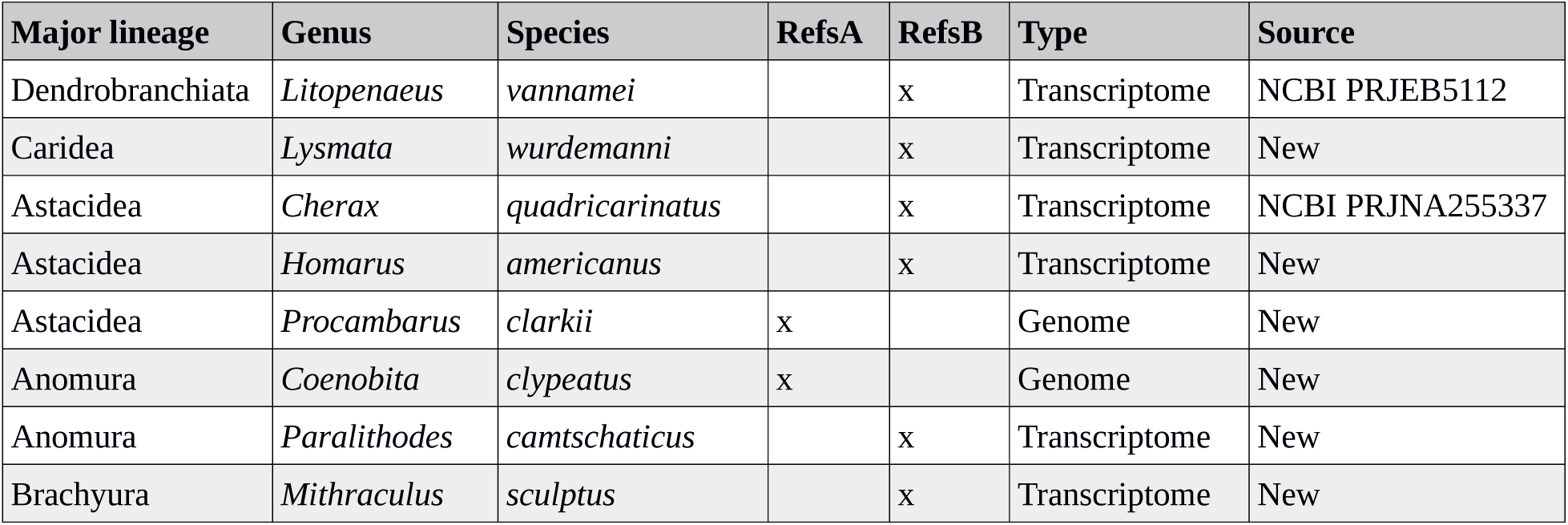
Genomes and transcriptomes used for preliminary probe design.

We used additional genomic resources (**Table S1**) to build alignments from six major lineages representing the diversity of decapods (Achelata, Anomura, Astacidea, Brachyura, Caridea, Dendrobranchiata). Raw reads from these species were mapped to the references above and used to extend probes into flanking regions [29]. For each combination of locus and major lineage, an alignment containing all recovered sequences (and lineage-specific reference sequence) was created in MAFFT. As each alignment contained at least one sequence derived from a genome, we were able to identify and mask intronic and repetitive regions, the latter identified based on the best-matching genomic region in the published red cherry shrimp (*Neocaridina denticulata*) genome [34]. Probes were tiled at 4x density across all sequences in each alignment and divided into two Agilent SureSelect XT kits (**Table S5**).

### (b) AHE sequencing and dataset assembly

From the Florida International Crustacean Collection (FICC), 89 species of decapods and seven additional crustaceans were selected for AHE sequencing. High molecular weight DNA was extracted from abdominal tissue, gills, or legs using the DNeasy Blood and Tissue Qiagen Kit following manufacturer’s protocol. A post-extraction RNase Treatment was performed on all samples to remove RNA contamination. AHE libraries were prepared from DNA extracts from 94 species (**Table S6**) at the FSU Center for Anchored Phylogenomics, following Lemmon et al. [27]. Libraries with 8 bp indices were combined in pools of 16 samples prior to enrichment with the Agilent SureSelect XT kits, then combined into two pools of 48 samples and sequenced in a single Illumina HiSeq2500 lane with 2x150 paired end reads, which were sanitized using Trim Galore! v0.4.0 [35]. Due to high divergence across Decapoda, we screened resulting AHE data for single-copy exons in the reference genome of the Chinese mitten crab, *Eriocheir sinensis* [36]. A total of 675 exons (L1-L675) were identified with ~40% coverage across AHE sequenced taxa. The *E. sinensis* amino acid sequences for these 675 loci were added to our data set and used as a reference locus set for Iterative Baited Assembly (IBA) and orthology screening following Breinholt et al. [31], except where noted in **Extended Methods 1f-g**.

### (c) Phylogenomics

The main data matrix used for phylogenetic analysis comprised 410 loci with at least 60% of the taxa represented in each locus (**Figure S2**). We analyzed both amino acid and nucleotide datasets, as nucleotides have been shown to support shallow relationships [37] and may be robust to differences among optimality criteria [38]. Analyzed matrices are summarized in **Table 2**. We inferred phylogenetic relationships using several methods, fully detailed in **Extended Methods 1i-j**. Bayesian inference was conducted with PhyloBayes v3.3f [39] using the site-heterogeneous CAT-GTR + G substitution model (only used for amino acid matrices). Maximum-likelihood analyses used IQ-TREE v1.6.3 [40] on 149 best-fitting partitions identified by PartitionFinder [41]. Coalescent (‘species tree’) methods were applied to investigate the role of incomplete lineage sorting, with maximum-likelihood gene trees inferred in IQ-TREE as inputs to estimate the species tree in ASTRAL-III v5.6.1 [42].

**Table 2.** Data matrix statistics.

| **Matrix** | **# loci** | **# amino acids** | **# base pairs** | **Average locus length (aa or bp, respectively)** | **% informative sites** | **% missing data** | **% GC content** |
| --- | --- | --- | --- | --- | --- | --- | --- |
| Amino acid | 410 | 28774 | n/a | 70 | 27 | 20 | n/a |
| Amino acid Dayhoff-6 recoding | 410 | 28774 | n/a | 70 | 14 | 20 | n/a |
| Amino acid top 50 | 50 | 5994 | n/a | 120 | 39 | 25 | n/a |
| Nucleotide | 410 | n/a | 86322 | 210 | 48 | 20 | 48.6 |
| Nucleotide positions 1+2 | 410 | n/a | 57548 | 140 | 24 | 20 | 45.8 |
| Nucleotide Degen recoding | 410 | n/a | 86322 | 210 | 11 | 20 | 45.2 |
| Nucleotide Degen positions 1+2 | 410 | n/a | 57548 | 140 | 17 | 20 | 43.8 |

### (d) Divergence time estimation

We identified 19 fossil calibrations across Decapoda, justified based on best practices ([43,44]; details in **Extended Data 2** and **Table S7**). All internal calibrations used soft bounds with 5% of the probability distribution allowed outside of the input ages, defined by a birth-death tree prior. We applied a gamma-distributed root prior based on crown Eumalacostraca [44] with a mean age of 440 Ma and SD 20 Myr. Divergence times were estimated in PhyloBayes using a fixed topology from our preferred tree (Bayesian CAT-GTR + G; discussed below), the CAT-GTR + G substitution model, multiple clock models, and two runs of four MCMC chains each.

## 3. Results and Discussion

### (a) Target capture success

We successfully sequenced targeted regions from 94 species representing 11 of 12 major decapod lineages (raw reads in NCBI BioProject XXX, assemblies in Dryad). We attempted to include *Neoglyphea inopinata*, one of only two living members of Glypheidea (deep sea lobsters with a diverse fossil record); however, multiple attempts to extract DNA from the limited tissue available to us did not render high-quality genomic extractions and failed during the probe capture (however, see [23] for mitogenome data). All other taxa we sequenced were quite successful, producing an average of 3,299,141 reads, with an average of 332 loci across samples that ranged from 57-405 assembled loci (**Figure S2**, **Table S8**). The final 410 loci ranged from 66-1683 bp with a total alignment length of 86,322 bp (**Table 2**). Taxa represented by >350 loci came from all major decapod lineages except Procarididea, demonstrating the efficacy of our probes across the entire clade. Using our enrichment kit, it will thus be possible for the community to easily sequence the same loci for large-scale phylogenomics spanning any decapods of interest.

The majority of nodes were congruent across different analyses, albeit with different levels of support (**Figure 2**), demonstrating that our large dataset is mostly cohesive and can resolve deep splits. We use the results from Bayesian inference with the CAT-GTR + G amino acid substitution model as the ‘best’ topology (**Figure 2**, first support square). This topology does not precisely match any previous result from morphological, Sanger, and/or mitogenomic data [19,21–23,25]. We include this tree over the Bayesian recoded topology because it had more nodes resolved, with higher support; the ways in which model misspecification drive contradicting topologies between these methods is still not understood [45]. Nucleotide analyses were not preferred because of both saturation in our data (**Figure S3**) and disagreement among results of different analyses [38].

**Figure 2.**
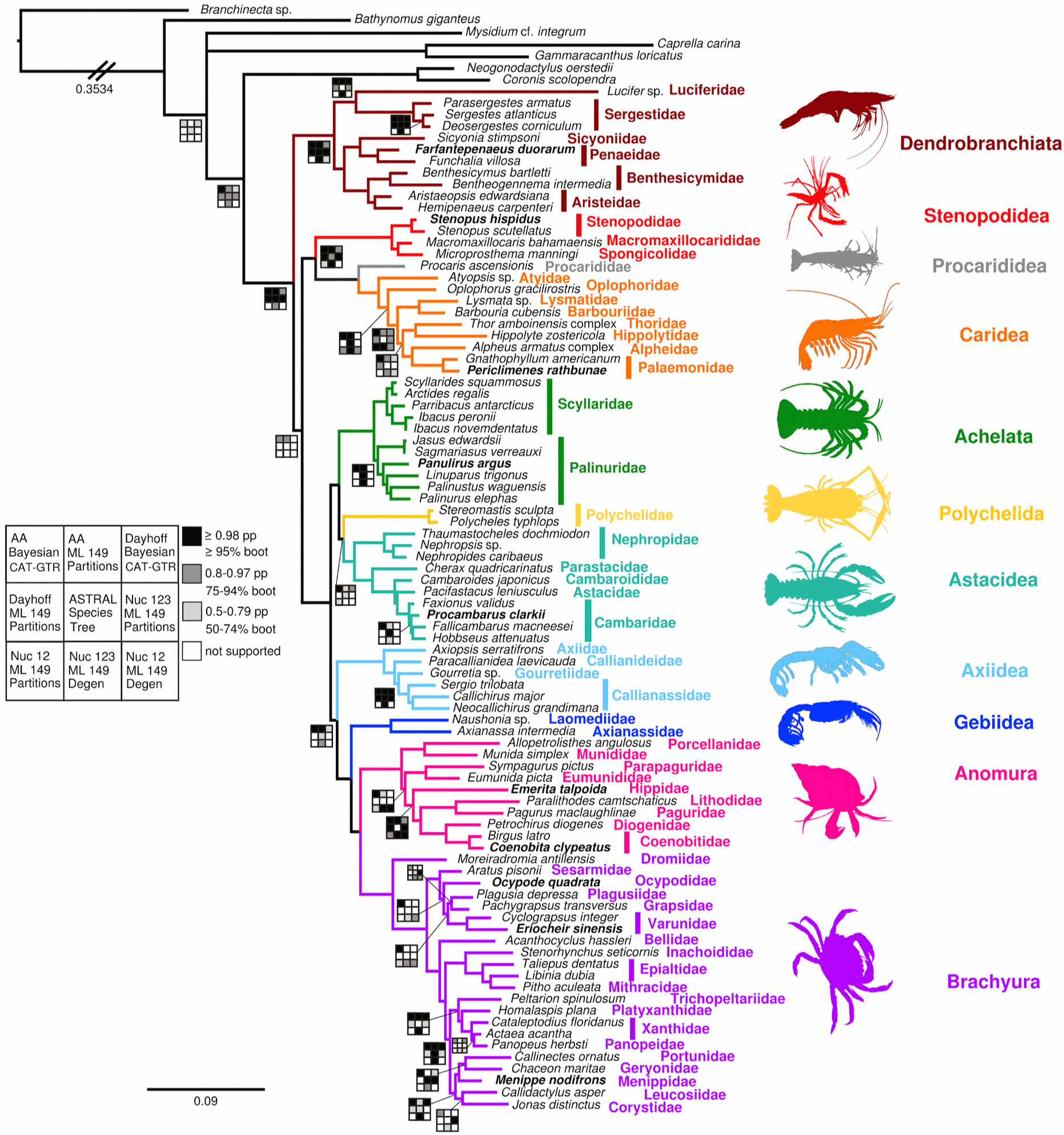
Phylogenetic hypothesis for Decapoda based on the topology from the Bayesian CAT-GTR + G analysis. Unlabeled nodes are considered strongly supported. Nodes where at least one analysis rejected the depicted topology are illustrated with rug plots showing the support values from each analysis. In rug plots, the illustrated topology is first row, first column. All alternative topologies are available in Dryad. Species used for probe design by shotgun whole genome sequencing in bold text. For clarity, the branch leading to the outgroup *Branchinecta* sp. (Anostraca) has been shortened, and the real length is indicated. Organism silhouettes are from PhyloPic (phylopic.org) or created by J.M. Wolfe.

### (b) Deep evolutionary history of decapods

Monophyly of Decapoda is supported in amino acid analyses (**Figure 2**); some nucleotide results find *Lucifer* (an apomorphic, epipelagic dendrobranchiate shrimp; **Figure 1a**) experiencing long branch attraction toward various outgroups. The most classical division in decapods, between suborders Dendrobranchiata (most food shrimp/prawns) and Pleocyemata (all other decapods), is supported by the unrecoded amino acid matrices (pp = 0.97/bootstrap = 93%), and contradicted by all others. The alternative hypothesis recovered is the natant (shrimp-like) decapods, with Dendrobranchiata, Stenopodidea, Procarididea, and Caridea forming a (usually poorly supported, ~50%) clade. We tentatively agree with the results supporting Dendrobranchiata and Pleocyemata, similar to the transcriptomic results of Schwentner et al. [26]. The polarity of one of the major characters separating these two clades, the lecithotrophic free-living nauplius larva in dendrobranchiates (as opposed to the egg-nauplius in pleocyemates), depends on whether Euphausiacea (krill) are most closely related to decapods, which we did not test. If euphausiids are the sister group of decapods, then pleocyemates have lost the free-living nauplius [46,47]; otherwise, the free-living nauplius is convergent in euphausiids and dendrobranchiates [48].

Within Pleocyemata, all infraorders receive full support for their monophyly (**Figure 2**). A single origin of the reptant or ‘crawling/walking’ decapods (Achelata, Polychelida, Astacidea, Axiidea, Gebiidea, Anomura, and Brachyura) is strongly supported. Numerous morphological characters have previously suggested monophyly of the reptant clade, e.g. a dorsoventrally flattened pleon, calcified body, anterior articulation of the mandibles formed by an elongated process of the molar region extending dorsally from the palp, anteroposterior rotation of walking legs, a short first pleomere, and spermatozoa with at least three nuclear arms [7]. Monophyly of reptant decapods is concordant with the majority of previous results [25], and almost certainly includes Glypheidea in addition [23].

Our posterior age estimate for the root of crown Decapoda (mean in the Late Ordovician at 455 Ma, 95% CI 512-412 Ma; **Figure 3**), was substantially older than most previous estimates [6,10,12,14,49], which largely fixed crown decapods in the Devonian. Our data include non-decapod outgroups, and the more crownward position of the Devonian calibration fossil *Palaeopalaemon newberryi* within Reptantia [44,50], resulting in older ages for deeper nodes.

**Figure 3.**
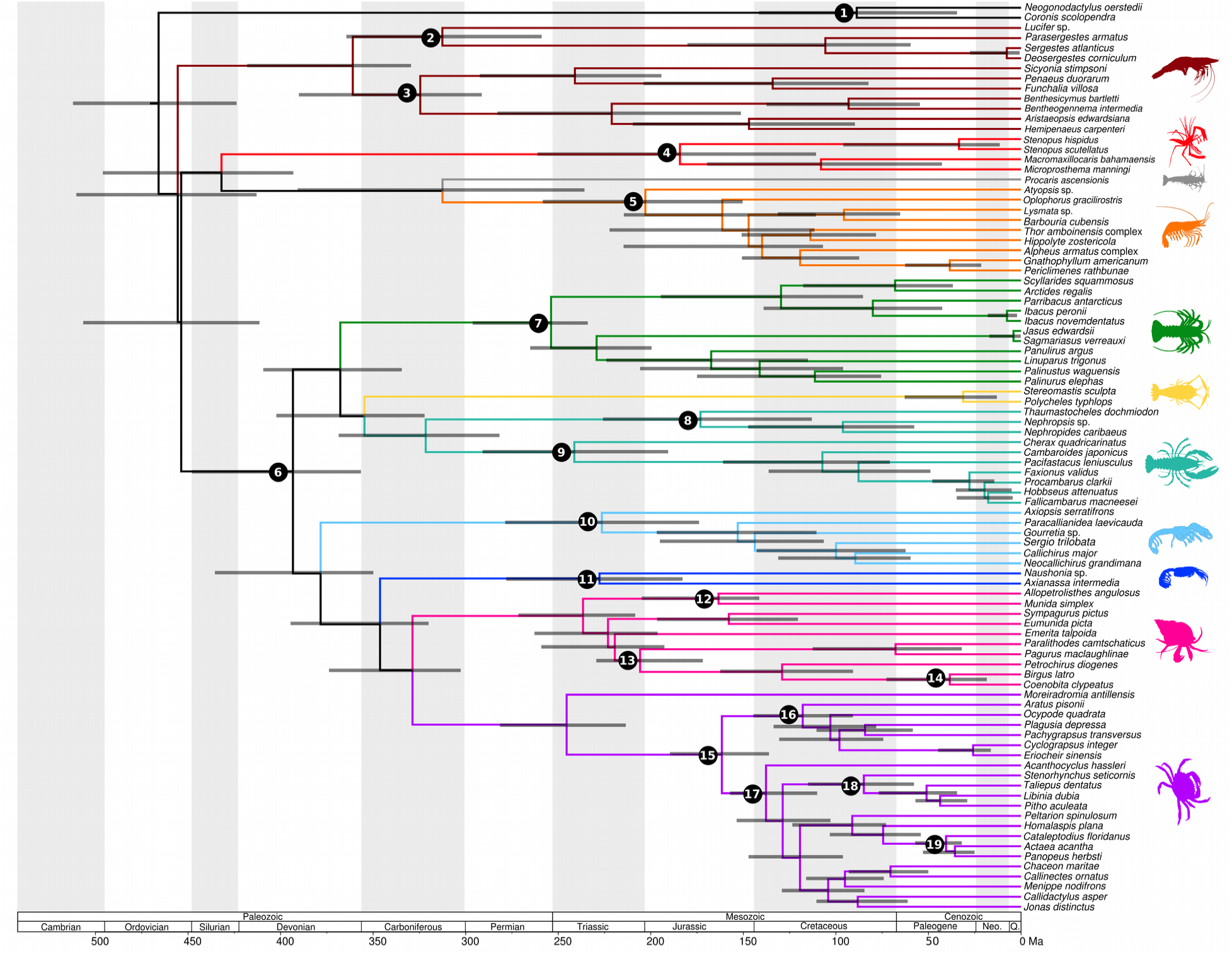
Divergence time estimates for Decapoda based on the topology in **Figure 2**. Posterior ages were estimated in PhyloBayes using the CAT-GTR + G substitution model, the CIR clock model, and a gamma distributed root prior of 440 Ma ± 20 Myr. Horizontal shaded bars represent 95% confidence intervals. Numbered circles represent nodes with fossil calibrations.

### (c) Evolutionary history of shrimp

Dendrobranchiate relationships are consistent (**Figure 2**), except the aforementioned long branch attraction of *Lucifer* to outgroups in the ASTRAL and nucleotide analyses. Amino acid results place this perplexing decapod with Sergestidae (pelagic shrimp), as suggested by morphological analysis, especially spermatophore morphology [51,52]. Crown dendrobranchiates diverged in the Late Devonian (**Figure 3**), with the two main clades Sergestoidea and Penaeoidea both diverging in the Pennsylvanian (about 73 Myr prior to the estimates of Ma et al. [53], and over 100 Myr older than the minimum ages suggested by Robalino et al. [54]). Although our mean posterior age of crown Penaeidae (134 Ma) is younger than the phylogenetically justified Late Jurassic crown fossil *Antrimpos speciosus* [54], which we did not use as a calibration, our 95% CI encompasses the fossil age of 151 Ma.

Among the pleocyemate shrimps, a sister group relationship between Stenopodidea (cleaner/boxer shrimp) and (Procarididea + Caridea) is strongly supported (**Figure 2**); similar topologies have been found in previous molecular and morphological analyses [8,11,26]. The sister group relationship of Procarididea (anchialine shrimp) and Caridea is supported by four-gene molecular analyses, the extended second pleurite overlapping the first and third somites, phyllobranchiate gills, and the form of the telson and uropods [6,55,56].

Within Caridea, we sampled eight families, compared with maximum 27 families in previous studies [18,57]. Nevertheless, our results are broadly concordant within the limits of the taxa we sequenced, producing a strongly supported backbone topology upon which future studies can build. The deepest split within carideans was between Atyidae (freshwater shrimp) and all others (**Figure 2**). Support for an alternative deep split of Atyidae and Oplophoridae from all other carideans (full nucleotide analyses) was weak, or a polytomy (recoded analyses). Our preferred topology agrees with previous work [18,57,58], although we are missing some deeper-diverging families. We support previous findings that the traditional concept of ‘Alpheoidea’ (snapping shrimp and allies) is not monophyletic and contains Palaemonidae (~200 genera; [18,57]). This larger Alpheioidea + Palaemonidae clade contains the majority of caridean diversity, possibly including Amphionidacea [19]. We inferred younger posterior ages (**Figure 3**) for the Procaridea-Caridea split (Pennsylvanian), and crown Caridea (Late Triassic), compared with previous analyses [6,58].

### (d) Evolutionary history of lobsters

Few previous analyses (from six or fewer nuclear genes: [11,16,17]) have recovered monophyly of the overall lobster body plan, which receives full support in all our analyses (**Figure 2**). The relationships among lobster infraorders, however, are poorly resolved. Amino acid analyses suggest a sister group relationship between Polychelida (blind deep sea lobsters) and Astacidea (clawed lobsters and crayfish), but with <70% support, except the Dayhoff recoded Bayesian analysis (pp = 0.86). Recent mitogenomic work shows Glypheidea potentially intervening as the sister group of Polychelida [23,24] within a paraphyletic lobster grouping. It is difficult to predict how the addition of Glypheidea would affect our topology, as it is also possible that the mitochondrial grouping results from long branch attraction (both Glypheidea and Polychelida have species-poor crown groups and morphologically diverse stem groups that predate their crowns by >150 Myr; [15,59]).

Achelata, with the crown members united by their unique phyllosomal larval stage, are monophyletic (**Figure 2**). Of the two constituent families, Scyllaridae (slipper lobsters) are monophyletic in all analyses. Palinuridae (spiny lobsters), however, are paraphyletic in amino acid recoded analyses, and in some nucleotide analyses. Based on monophyletic Palinuridae (**Figure 3**), their crown divergence occurred in the Late Triassic (similar to previous estimates; [15]), and crown Scyllaridae in the Early Cretaceous (about 87 Myr younger than Bracken-Grissom et al. [15]). These age estimates predate the wealth of Jurassic and Cretaceous fossil achelatan larvae [60,61], implying bizarre stem-groups may have persisted throughout the Mesozoic alongside the crown. Our phylogenetic results also support the division of Palinuridae into distinct clades of Silentes and Stridentes, the latter bearing an enlarged antennular plate used in sound production in adults [15,62,63]. Indeed, the palinurid genera that are close to Scyllaridae in these alternative analyses are the included members of Silentes (*Jasus* and *Sagmariasus*), further supporting the auditory behavior of Stridentes as a clade synapomorphy, having diversified in the Jurassic.

Relationships within Astacidea were similar to results from a combined Sanger sequencing and taxonomic synthesis approach [64,65]. Crown Astacidea diverged in the Pennsylvanian, and crown Nephropidae (‘true’ lobsters) diverged in the Early Jurassic (**Figure 3**). The split between southern hemisphere crayfish (Parastacidae) and northern hemisphere crayfish occurred in the Middle Triassic around 241 Ma, prior to the breakup of Pangaea [15].

### (e) Evolutionary history of mud/ghost shrimp

The mud/ghost shrimps Axiidea and Gebiidea (formerly Thalassinidea) are not monophyletic, as has been shown in previous molecular work [66]. Most of our amino acid analyses produce a paraphyletic mud shrimp group (**Figure 2**), with limited but clear support for Axiidea as the sister group to the Gebiidea and Meiura clade (i.e. the Monochélie of de Saint Laurent [67], which is strongly supported herein). There is precedent for our mud shrimp and crab clade based on morphology (where thalassinidean monophyly is assumed [7]), Sanger results [14], and on some mitogenomic analyses [20,22,23]. The alternative hypothesis we recovered (in most of the nucleotide analyses) is mud shrimp polyphyly, with Axiidea as sister group to the lobster clade. Loss of chelae on pereiopods posterior to the first was previously proposed asconvergent in Meiura and members of Gebiidea [7]; our topology suggests this character is the eponymic synapomorphy of the Monochélie [66,67]. Our divergence time analysis suggests that the mud shrimp + crab clade diverged in the Late Devonian (**Figure 3**), with crown Monochélie diverging in the Mississippian, and both crown Axiidea and crown Gebiidea in the Late Triassic. These posterior age estimates are older than previous studies [14] or a literal interpretation of the fossil record [68].

### (f) Evolutionary history of crabs

Meiura, the monophyletic relationship between Anomura (‘false’ crabs) and Brachyura (true crabs), is strongly supported (**Figure 2**). This is important because a number of Sanger analyses [6,10,13,16,19] purported to refute meiuran monophyly. Several synapomorphies have been proposed, such as a short asymmetric flagella on the antennule, bent exopods on the maxillipeds, and fusion of ganglia borne on the first pleomere and thoracic mass [7]. Carcinization, the overall crab-like body plan including a flattened carapace with lateral margins, fused sternites, and strongly bent abdomen [69] has been suggested as a developmentally co-opted trait of Meiura with a ‘tendency’ to evolve repeatedly [70,71]. Our topology suggests at least four separately carcinized clades in Anomura, and one in Brachyura; however, increased taxon sampling will complicate character distribution [14,69] and introduce secondary losses, such as in frog crabs [72,73].

Within Anomura, we recover support for Hippidae (mole crabs) as the sister group of ‘Paguroidea’ (king crabs and most hermit crabs; **Figure 2**) rather than the outgroup to all other anomurans [14,74]. Precedent for a sister group relationship of Hippidae to Paguroidea comes from mitochondrial gene rearrangements [71], from gross adult morphology (e.g. shape of carapace and sternites [75]), and from characters of the foregut ossicles [76]. Note, as in past molecular-only results [14,74], Parapaguridae are more closely related to Eumunididae (squat lobsters), rendering hermit crabs polyphyletic with potentially convergent evolution of asymmetrical abdomens [74]. Recent mitogenome research [24] displayed dramatically different relationships among Anomura, including a different topology for hermit crab polyphyly, but mitochondrial data are best for relationships within families, and weaker for deep splits as they represent a single locus: long established as a weak approach to phylogenetic estimation. Posterior divergence estimates (**Figure 3**) from crown Anomura to crown Paguroidea span a narrow interval of about 22 Myr in the Late Triassic, with each node about 20 Myr older than previous estimates [14]. Note that we recover these posterior ages based on only soft maximum priors (i.e. not minima) from the Late Triassic *Platykotta akaina* [14,77], as it may be placed outside the meiuran crown-group [78]. We also observe a conflicting split between Galatheoida and all other anomurans (**Figure 2**), where preferred analyses support previous molecular-only results [14,74]. The alternative, a clade of monophyletic squat lobsters, porcelain crabs, and Parapaguridae, is supported in recoded amino acid, ASTRAL, and non-recoded nucleotide trees. These relationships could be clarified by sampling additional squat lobster and hermit crab groups.

Our analyses strongly support the traditional morphological divisions within Brachyura (**Figure 2**), with podotremes (represented by Dromiidae, or sponge crabs; gonopores located on the pereiopod coxa) as the deepest split in the Late Triassic (**Figure 3**), and eubrachyurans divided into Thoracotremata (gonopores located on the sternum) and Heterotremata (female gonopores located on the sternum, male gonopores on the coxa). Each of the two eubrachyuran branches diverged in the Early Cretaceous, with diversification among families mainly in the Late Cretaceous. Within Thoracotremata, all our results reject monophyletic Grapsoidea (**Figure 2**). The focal tree supports Sesarmidae as the outgroup to other families, but other analyses support either Plagusiidae or Varunidae. Within Heterotremata, we recover support for several clades that have been previously defined [12], at least within our taxon sampling: Majoidea (spider and decorator crabs: Epialtidae, Inachoididae, and Mithracidae), Xanthoidea (mud crabs: Panopeidae and Xanthidae), and Portunoidea (swimming crabs: Portunidae and Geryonidae). Within Majoidea, the family Epialtidae is paraphyletic with respect to Mithracidae, suggesting continued evaluation of larval morphology [12,79,80]. This is the best supported region of the heterotreme tree. As we only sampled 19 of 96 total brachyuran families, it is unsurprising that our analyses conflicted for remaining clades. Important AHE target taxa for improved resolution include Raninidae, Cyclodorippidae, and Homolidae (all podotremes), Gecarcinidae and Pinnotheridae (pea crabs, within thoracotremes), primary freshwater heterotremes [12], and xanthoid relatives [81].

### (g) Divergence times

We present the posterior age results from divergence time analysis using the CIR autocorrelated clock model (**Figure 3**), as it is more biologically realistic [82]. Unlike in broader studies of arthropods [83] and myriapods [84], posterior credibility intervals were similar for many nodes regardless of which clock model we applied (**Figure S4**). Although we only used the top 50 loci as sequence data to estimate divergence times [85], the posteriors hewing close to the effective prior are not necessarily problematic (e.g. [86–88]). Similarities between effective prior and posterior distributions are also present for nodes we did not explicitly calibrate (**Figure S4b**), though they are free to vary according to the birth-death tree prior. This effect is less pronounced for non-reptantian nodes, which have scant fossil information and essentially uniform maxima as described by Brown & Smith [89].

Overall, our divergence time estimates imply a significant cryptic history for decapods, encompassing most of the Paleozoic (**Figure 3**). Perhaps our results will motivate revision of Paleozoic fossils that have been suggested as decapods, such as *Imocaris* spp. [78,90,91], angustidontids [92,93], or poorly constrained natantian shrimp [94], in a more explicit phylogenetic framework. We also infer a lack of cladogenesis among the deep lineages during the Permian, followed by diversification in most crown groups in the Triassic. Although we did not explicitly calibrate most nodes using Triassic fossils, and molecular data alone cannot accurately estimate diversification [95], it is striking that our divergence time analysis infers appearance of the modern decapod clades following the largest known mass extinction 251 Ma, replacing and innovating ecological roles as important members of the Modern evolutionary fauna [96–98]. Moreover, the most species-rich lineages, Caridea, Anomura, and Brachyura, each show deep divergences during the Jurassic and family-level diversification in the Cretaceous, concurrent with the radiation of modern reef-building corals [99], a major habitat and source of biodiversity for these crustaceans [100,101].

## 4. Conclusion

Our well-resolved dated phylogeny may inform comparative evolutionary topics, such as the evolution of visual systems in deep sea and cave environments [64,102], evolution of major body plan features [14,69,103], the role of symbiosis [104–106], evolution of behavior [107], macroevolutionary trends in physiology and habitat through time [100,108], conservation biology and vulnerability to climate change [49,109,110], and more. The new enrichment kit we have generated will permit an inexpensive expansion of taxon sampling across Decapoda, via our large-scale matrix of loci conserved across 450 Myr, to accelerate discoveries in a fascinating invertebrate clade.

## Data accessibility

Extended Methods and Fossil Calibrations, Figures S1-S4, and Tables S1-S8 are available as Electronic Supplementary Material. Raw reads are available in the NCBI BioProject: XXX. Assemblies, matrices, and resulting tree files are available at the Dryad Digital Repository, provisional link: https://datadryad.org/review?doi=doi:10.5061/dryad.k7505mn. Scripts for this paper are available at https://github.com/jessebreinholt/proteinIBA.git.

## Authors’ contributions

H.D.B.G. and K.A.C. conceived the project, J.M.W., K.A.C., M.E.S., and H.D.B.G. acquired samples, L.E.T. and H.D.B.G. extracted DNA, J.M.W. and M.E.S. extracted RNA, A.R.L. and E.M.L. developed probes and conducted sequencing, J.W.B. developed, assembled, screened orthologs and processed AHE data into matrices, J.W.B. and J.M.W. conducted phylogenetic analysis, J.M.W. vetted fossil constraints, performed divergence time analyses, created figures, and wrote the manuscript with input from all authors.

## Acknowledgments

We thank Mercer Brugler, Jorge Perez-Moreno, Shaina Simon, and Juliet Wong for assistance with extractions, and Michelle Kortyna, Sean Holland, and Ameer Jalal at the FSU Center for Anchored Phylogenomics for assistance with probe design and data collection. We acknowledge use of the Engaging Cluster at MGHPCC. This is contribution #X for the Center for Coastal Oceans Research in the Institute for Water and Environment at Florida International University.

## Extended Data

### 1. Extended Methods

#### (a) Whole genome sequencing and QC

Whole genome sequencing (WGS) was conducted for nine exemplar decapod species (**Table S2**): *Panulirus argus* (Achelata: Palinuridae), *Coenobita clypeatus* (Anomura: Coenobitidae), *Emerita talpoida* (Anomura: Hippidae), *Procambarus clarkii* (Astacidea: Cambaridae), *Menippe nodifrons* (Brachyura: Menippidae), *Ocypode quadrata* (Brachyura: Ocypodidae), *Periclimenes rathbunae* (Caridea: Palaemonidae), *Penaeus duorarum* (Dendrobranchiata: Penaeidae), and *Stenopus hispidus* (Stenopodidea: Stenopodidae). These species were targeted as they span the phylogenetic breath of Decapoda, and could be freshly collected (or represented recently collected material). All material was selected from the Florida International Crustacean Collection (FICC) or newly collected in localities around southern Florida. All identifications were done by H. Bracken-Grissom using dichotomous keys for these groups.

DNA was extracted from gills, legs or abdominal muscle tissue using the Qiagen Blood and Tissue Extraction Kit following manufacturers protocols. For these 9 decapod lineages, low-coverage genome data were newly collected after DNA extracts were sonicated to a distribution of 200-500 bp and used to produce 8 bp single-indexed Illumina libraries following Lemmon et al. [1] and Prum et al. [2]. Libraries were pooled and sequenced on 18 PE-150 Illumina HiSeq2500 lanes with onboard clustering, producing 711 Gb of data (3x to 16x genomic coverage). After sequencing, paired reads passing Illumina Casava High Chastity filter were merged following Rokyta et al. [3], during which sequencing errors were corrected and library adapters were removed. These genomes were not assembled; the short reads were screened for probe design (discussed below).

#### (b) Transcriptome sequencing

Transcriptomes were sequenced from multiple developmental stages for each of four exemplar decapod species: *Homarus americanus* (Astacidea: Nephropidae), *Lysmata wurdemannii* (Caridea: Lysmatidae), *Mithraculus sculptus* (Brachyura: Mithracidae), and *Paralithodes camtschaticus* (Anomura: Lithodidae). For adult females, brain and/or muscle tissues were dissected. For embryos (mid-germband stage) and larvae (first zoea), several individuals were pooled into 1-3 replicates for each stage. Each sample was preserved in RNALater for transport to AMNH. Collecting and sample information are found in **Table S3**.

Total RNA was extracted separately from individual tissues. Prior to immersion in RNALater, whole animals were rinsed in 10% bleach followed by deionized water (to minimize microbial contaminants). Samples were separately macerated in Trizol using the Xpedition mechanical sample processor (Zymo Research), followed by RNA extraction with the Direct-zol kit (Zymo), including a poly-A binding step to isolate mRNA. The quality and quantity of mRNA was measured with the BioAnalyzer 2100 (Agilent) and Qubit 2.0 fluorometer (Invitrogen). cDNA libraries were generated with TruSeq DNA preps (Illumina) at Weill Cornell Medical College (for *L. wurdemanni*) and the New York Genome Center (for the other three species). Transcriptomes were sequenced at the above respective labs, using Illumina HiSeq with 2×100 paired end reads, and 9-10 samples multiplexed per flow cell.

#### (c) Transcriptome QC and assembly

Raw Illumina reads were filtered for each sample using a Galaxy workflow based on the FASTQ toolkit, following [4]. Briefly, adapter sequences were removed, followed by FASTQ Groomer, removal of sequencing artifacts, and quality filtering. *De novo* transcriptome assembly of filtered reads, combining all sequenced samples of a species, used Trinity [5] with the default minimum k-mer threshold abundance (=1, so all k-mers were used in assembly). As in Alexandrou et al. [4], Trinity output was re-assembled using iAssembler [6]. The number of raw and filtered reads and assembly statistics are compiled in **Table S4**.

#### (d) Probe design

Target AHE loci were identified at the FSU Center for Anchored Phylogenomics (www.anchoredphylogeny.com). using the approach successfully employed for other arthropod groups, including Diptera [7], Arachnida [8], Hemiptera [9], Coleoptera [10], Neuroptera [11], and Lepidoptera [12]. Genomic resources from 23 species representing a variety of Decapoda were obtained from published and unpublished sources (see **Table S1** for details), including the nine whole genomes and four transcriptomes described above.

To identify AHE loci in Decapoda, genomic reads from two species (RefsA in **Table 1**) identified in preliminary tests to yield the highest locus recovery, *Coenobita clypeatus* (Anomura: Coenobitidae) and *Procambarus clarkii* (Astacidea: Cambaridae) were mapped to reference sequences from *Tribolium castanaeum.* The *Tribolium* sequences were obtained from the Coleoptera AHE probe design alignments developed by Haddad et al. [10], who targeted protein coding regions conserved across Insecta. The mapping process, which follows that described in detail by Hamilton et al. [8], identifies matching reads using a 17 bp of 20 bp spaced-k-mer threshold for preliminary matching and 55% similarity score for final placement of a read. For each locus, reads matched by these criteria were aligned and then extended into flanking regions (see [8] for details). For each locus, the obtained decapod sequences were aligned together with the corresponding *Tribolium* sequence using MAFFT v7.023 [13]. Geneious R9 (Biomatters Ltd.; [14]) was used to identify and select well-aligned regions that were utilized downstream. This process resulted in 823 preliminary AHE target sequences.

To develop a suitable reference for each of the six selected decapod ‘major lineages’ (Achelata, Anomura, Astacidea, Brachyura, Caridea, and Dendrobranchiata), we scanned for AHE loci in six assembled transcriptomes (RefsB in **Table 1**); these include the four newly sequenced transcriptomes above, as well as *Litopenaeus vannamei* (Dendrobranchiata: Penaeidae) and *Cherax quadricarinatus* (Astacidea: Parastacidae [15]). The references we used were the 823 sequences from *C. clypeatus* and *P. clarkii* that are described above. Following Hamilton et al. [8], we identified for each locus the best-matching transcript in each of the six transcriptomes. The resulting sequences were aligned using MAFFT [13] and inspected in Geneious. After selecting the well-aligned regions and removing 21 poorly aligned sequences, loci represented by fewer than four of the six selected major lineages were removed from downstream analyses. Removal of overlapping loci (identified as having one or more shared 60-mer for any species) resulted in 352 final target loci.

The remaining genomic resources were then utilized to build alignments representing the diversity within each of the six selected major lineages. Raw reads from 16 additional species (see **Table S1**) were mapped to major lineage-specific reference sequences (developed above) and used to extend the resulting sequences into flanking regions [8]. The best matching genomic region in the assembled *Neocardina denticulata* genome (Caridea: Atyidae; [16]) was also identified (4000 bp containing each region was utilized downstream). For each locus × major lineage combination, an alignment containing all of the recovered sequences (together with the major lineage-specific reference sequence) was then generated using MAFFT and inspected in Geneious. As each alignment contained at least one sequence derived from a genome, we were able to identify and mask intronic regions. We also masked repetitive regions identified using k-mer counts in the *N. denticulata* genome (see Hamilton et al. [8] methodological details). Probes were tiled at 4x density across all sequences in each of the alignments, and similar probes were removed. **Table S5** provides details of the target size and number of taxa representing each major lineage. The probes were divided across two Agilent SureSelect XT kits as follows: 1) Decapoda1a ELID=801331 containing Den1, Car1, and Ast1 (56698 probes total), and 2) Decapoda1b ELID=801341 containing Ach1, Ano1, and Bra1 (54854 probes total).

#### (e) AHE sequencing

Samples of 94 species (**Table S6**) were processed at the FSU Center for Anchored Phylogeny (www.anchoredphylogeny.com) following Lemmon et al. [1] and Prum et al. [2]. Briefly, DNA extracts passing QC by Qubit were sonicated to a size range of 200-500 bp and prepared to include single 8 bp library adapters. Libraries were pooled in groups of 16 and enriched using the aforementioned probes (probes from the two kits were pooled prior to the enrichment reaction). Enriched library pools were then pooled for sequencing such that each of two 48-sample pools were sequenced on an Illumina HiSeq2500 lane with a PE150 protocol and 8 bp indexing (85 Gb total).

#### (f) AHE QC and assembly

Paired-end raw Illumina reads were cleaned using Trim Galore! v0.4.0 [17], allowing a minimum read size of 30 bp and trimming to remove bases with a Phred score below 20. To build a reference set of loci for Iterative Baited Assembly (IBA; [12]), we first identified single-copy exons in the published *Eriocheir sinensis* genome (Brachyura: Varunidae; [18]; NCBI PRJNA305216) using single-hit and ortholog location genome mapping following Breinholt et al. ([12]; scripts: s_ hit_checker.py and ortholog_filter.py). These single-copy exons from *E. sinensis* were screened against *de novo* Bridger assemblies of our cleaned AHE data (using default parameters: [19]) using script genome_getprobe_TBLASTX.py (available: https://github.com/jessebreinholt/proteinIBA.git: [20]) that uses tblastx instead of blastn and ortholog_filter.py [12]. Loci that had 40% or more of the taxa sequenced for AHE samples in this study were used as a 675 locus reference set called crab_ref1, which were translated into amino acids as bait for IBA. The IBA successively uses USEARCH v7.0 [21] and Bridger v2014-12-01 [19] to assemble each locus in the reference set, and ensure it hits the targeted probe of the reference taxon (*E. sinensis*). IBA was altered to take an amino acid reference using the script protein_IBA.py (available: https://github.com/jessebreinholt/proteinIBA.git). For protein_IBA, the k-mer size was set to 25 and k-mer coverage of assembled sequences was set at 10x.

#### (g) Alignment, orthology, and alignment trimming

The blast table and the assembled sequences output from IBA for each locus were processed with protein_dir_fixer.py (available: https://github.com/jessebreinholt/proteinIBA.git). This script put all sequences in the correct direction and trimmed them to match the probe region for ortholog screening. The sequences were queried using tblastx against the *E. sinensis* genome, and screened for orthology using the ortholog_filter.py following Breinholt et al. [12]. To control for possible contamination, we selected the sequences for every taxon with the lowest ‘comp’ number output by IBA, which is the set of sequences and isoforms that received the most reads and depth. This method provided results comparable to the contamination filter and removing duplicates step [12] without having to make assumptions about contamination using sequence similarity based on taxonomy. The full-length sequences of the inferred orthologs were aligned with MAFFT v7.245 [13]. As very little data appeared to be conserved or alignable across Decapoda in the introns outside the probe region, we trimmed to the probe region (i.e. exon) using the Extract_probe_region.py script [12]. Consensus of isoforms for each sequence were made to reduce to a single sequence per locus and taxon using FASconCAT-G v1.02 [22] following Breinholt et al. [12].

#### (h) Data matrix construction

The main data matrix comprised 410 loci with at least 60% of the taxa represented in each locus (average occupancy per locus = 81%; completeness score as the total number of unambiguous characters divided by the size of the alignment = 80%). We tested ALICUT/ALISCORE [23] to remove poorly aligned regions, but only three nucleotides were removed (and no amino acids), thus this step was not included in the final dataset. A heatmap of pairwise amino acid completeness was constructed in ALISTAT [24] for all targeted loci (**Figure S2**). Details of final amino acid and nucleotide matrices are described in **Table 2**.

#### (i) Systematic error

Systematic error is implicated in phylogenomics where the data violate assumptions of the model, leading to convergence of ever-larger datasets towards an incorrect topology with high support values (e.g. [25,26]). At the nucleotide level, both site-specific and lineage-specific biases can strongly influence phylogenetic results. We investigated site-specific biases (i.e. multiple substitutions at the same site, or nucleotide saturation) by first building a saturation plot for each codon position with DAMBE v6 [27]. The saturation plot (**Figure S3**) indicated strong saturation of the third codon position, i.e. that this position evolved at a faster rate. Therefore, we conducted analyses on the full nucleotide set, as well as codons 1+2 only [28] as a form of data removal.

Broader biases in nucleotide compositional heterogeneity (lineage- and/or site-specific) were investigated using datasets that were recoded to exclude synonymous substitutions. These substitutions may or may not be saturated; biological base compositional bias is also accounted for. Recoding used ambiguity codes in the degen v1.4 Perl script [12,29]. Degen recoding does not change the matrix size; it merely reduces the degrees of freedom for possible substitution combinations.

In the amino acid dataset, site-specific amino acid biases may be accounted for with the CAT-GTR substitution model, but lineage-specific compositional biases do not have easily implementable models [30]. Therefore we adopted a similar approach as we did for nucleotides, using Dayhoff-6 recoding [30–32]. As with degen recoding, it does not change the matrix size, but groups the 20 amino acids into six classes that are similar on the basis of their biochemical properties, reducing the number of possible substitution combinations. This recoding strategy has been recommended for codon usage biases observed in pancrustacean nuclear genes in particular [32], though it has also been observed to collapse nodal support in other taxa e.g. [30,33].

#### (j) Phylogenetic analysis

For maximum likelihood (ML) analysis, PartitionFinder [34] on the 410 locus dataset (restricted to all versions of the LG model, including free rate models) obtained 149 best-fitting partitions. The best-fitting substitution models for each of the 149 partitions were selected in IQ-TREE v1.63 [35]. In IQ-TREE, 50 independent searches were run with different random seeds. All topologies were the same in each search, though branch lengths and scores differed. Support was assessed with 250 nonparametric bootstraps, with *a posteriori* convergence determined in RaxML using the MR, MRE, and MRE_IGN criteria (majority rule; convergence at 100) and FC criterion (frequency-based; convergence at 150). ML analyses were conducted on all matrices described in **Table 2**, except amino acid top 50.

For Bayesian inference, we used two chains and the CAT-GTR + G site-heterogeneous substitution model implemented in PhyloBayes v3.3f [36]. All Bayesian analyses were conducted on amino acid matrices; one matrix was recoded into the Dayhoff-6 functional groups. An automatic stopping rule was implemented, with tests of convergence every 100 cycles, until the default criteria of effective sizes and parameter discrepancies between chains were met (50 and 0.3, respectively), and with the bpcomp and tracecomp commands. Trees and posterior probability distributions were then generated from completed chains after the initial 20% of sampled generations were discarded as burn-in.

Coalescent (‘species tree’) methods were applied to investigate the role of incomplete lineage sorting e.g. [37,38]. ASTRAL is statistically consistent with the multispecies coalescent, and can be effective if there are sufficient gene trees matching the ‘true’ tree [39]. As input, we used maximum likelihood gene trees calculated by IQ-TREE (only on amino acid data, otherwise as above) with SH-like support. The species tree was then estimated in ASTRAL-III v5.6.1 [40], collapsing nodes with <10% support, and estimating branch support using quartet node support (the percentage of quartets in gene trees that agree with a branch).

#### (k) Divergence time estimation

Divergence times were estimated in PhyloBayes v3.3f [36] using a fixed topology input based on the Bayesian CAT-GTR + G tree depicted in **Figure 2**. Preliminary runs indicated that the distant outgroups may exhibit heterotachy, so all outgroups were removed from this topology, except the two stomatopods. Due to the size of our data matrices and time to convergence, we used an amino acid alignment of only the top 50 loci (those with the most similar gene topologies to the concatenated fixed topology; [41]) in PhyloBayes under the CAT-GTR + G substitution model, and two runs of four chains. We compared the uncorrelated gamma multipliers (UGM) relaxed clock model [42], and the autocorrelated CIR model [43]. The root prior was defined based on the Eumalacostraca node [44], thus applying a gamma distribution with a mean of 440 Ma and SD of 20 Myr. All 19 internal fossil priors (justified in **Extended Data 2**) used soft bounds with 5% of the probability distribution allowed outside of the input ages, defined by a birth-death model [45]. An automatic stopping rule was implemented, with tests of convergence every 100 cycles, until the default criteria of effective sizes and parameter discrepancies between chains were met (50 and 0.3, respectively), and with the bpcomp and tracecomp commands. Trees and posterior probability distributions were then generated from completed chains after the initial 20% of sampled generations were discarded as burn-in. We also compared estimated posteriors to the truncated effective prior (by removing sequence data using the -prior flag in PhyloBayes; [46,47]; **Figure S4**).

### 2. Fossil Calibration Justifications

We used 19 fossil calibrations, following the criteria of Parham et al. [48]. Node numbers listed below correspond to labels in **Figure 3**. A summary of fossils and their ages is provided in **Table S7**.

1. ***Node.*** This node comprises crown Verunipeltata (‘Stomatopoda’, or mantis shrimp) in our tree, with nomenclature as previously discussed [44,49], and monophyly matching Van Der Wal et al. [50]. All calibration data as in Wolfe et al. [44], node 51.
2. ***Node.*** This node represents crown Sergestoidea. In our phylogeny, this is the clade comprising *Lucifer*, *Parasergestes*, *Sergestes*, and *Deosergestes*, their last common ancestor and all of its descendants. ***Fossil specimens.** Paleomattea deliciosa* Maisey and Carvalho 1995 [51]. Holotype AMNH (American Museum of Natural History) 44985, carapace and abdominal segments dissolved from acid prep of teleost fossil. ***Phylogenetic justification.*** The phylogenetic position of this fossil was determined in a total evidence analysis [52]. It was found within crown Sergestoidea in their taxon sampling, sister taxon to *Acetes.* As it was outside of *Sergia* + *Deosergestes*, and we did not sample *Acetes*, we allow *P. deliciosa* to calibrate the clade including *Lucifer.* Note that a likely member of Luciferidae was recently discovered from the same deposits [53], and could be appropriate for this node as well. ***Age justification.*** Minimum as in Wolfe et al. [44], node 43. As the oldest decapod, *Palaeopalaemon newberryi* (node 6), is crownward of shrimps and prawns, a phylogenetic bracketing approach to obtain a soft maximum age for these groups is not appropriate. If we were to use a maximum age from the oldest crown Malacostraca, which would be 434.2 Ma [44], it would create priors that come into conflict with that of *P. newberryi* and its older age range for all decapods. Thus we conservatively adopt a soft maximum age of 521 Ma, as in node 1.
3. ***Node.*** This node represents crown Penaeoidea (penaeid shrimp/prawns). In our phylogeny it is the clade comprising Penaeidae, Sicyoniidae, Aristeidae, and Benthesicymidae, their last common ancestor and all of its descendants. Monophyly of this clade was also supported by a recent total evidence (morphology + three nuclear gene) phylogeny [52]. ***Fossil specimens.** Ifasya madagascariensis* Van Straelen 1933 [54]. Several specimens examined for coding in Robalino et al. [52], including: MSNM (Museo di Storia Naturale di Milano, Milan, Italy) i11309, i9311, i9408, i14229, i9383, i9406, i9328, i11243, and i9328. ***Phylogenetic justification.*** The phylogenetic position of this fossil was determined in a total evidence analysis [52]. It was found within crown Penaeidae, thus also crown Penaeoidea. ***Age justification.** I. madagascariensis* was discovered in the Ambilobé region of northwest Magadascar [55,56]. The fossiliferous sediments bear the index conchostracan *Euestheria* (*Magniestheria*) *truempyi*, which may be correlated to the Bernburg Formation of Germany [57]. Although this correlation was used as an indicator of Olenekian age for Ambilobé [57], revisions to conchostracan biostratigraphy correlate *M. truempyi* to the Dienerian substage of the global Induan stage, Early Triassic [58]. The minimum age of the Induan (or the lower boundary of the Olenekian) is 251.2 Ma, based on the 2017 International Chronostratigraphic Chart, thus providing a minimum age for *I. madagascariensis.* Soft maximum as in node 2 herein.
4. ***Node.*** This node represents crown Stenopodidea (cleaner/boxer shrimp). In our phylogeny, this is the clade comprising the (currently recognized) genera *Stenopus*, *Microprosthema*, and *Macromaxillocaris*, their last common ancestor and all of its descendants. ***Fossil specimens.** Phoenice pasinii* Garassino 2001 [59]. Holotype, MSNM i24799. ***Phylogenetic justification.*** Recently, a need to revise the systematics of crown Stenopodidea has been identified based on a molecular phylogeny [60]. Only three fossil species have been described, and none have yet been evaluated using phylogenetics. Of these, two, *Devonostenopus pennsylvaniensis* Jones et al. 2014 [61] and *Jilinocaris chinensis* Schram et al. 2000 [62], were placed within Stenopodidea on characters that were admittedly poorly preserved and not conclusive. In contrast, *P. pasinii* is better preserved, and has a number of diagnostic characters confirming its stenopodidean affinity [59]. In particular, the subtriangular telson and presence of two longitudinal dorsal carinae on the uropodal endopods suggest a relationship to the Stenopodidae [59]. While it is not clear if *P. pasinii* belongs within crown versus stem Stenopodidae, it is reasonable to assign this fossil within crown Stenopodidea. ***Age justification.** P. pasinii* was discovered in the Hakel/Hâqel (holotype) and Hadjula/Hjoûla outcrops, northeast of Beirut, Lebanon [59]. Presence of the ammonite *Allocrioceras* cf. *annulatum* in the Hjoûla limestone correlates the deposit to the *Sciponoceras gracile* Zone in the Western Interior of the USA and the *Metoicoceras geslinianum* Zone globally [63]. The *S. gracile* Zone has a cyclostratigraphic minimum age of 94.39 Ma ± 0.12 Myr [64], providing a minimum age constraint of 94.27 Ma for *P. pasinii.* Soft maximum as in node 2 herein, to allow for the possibility that *Devonostenopus* is within the crown.
5. ***Node.*** This node represents crown Caridea. In our phylogeny, this clade is represented by Alpheidae (snapping shrimp), Atyidae (freshwater shrimp), Barbouriidae, Hippolytidae (cleaner and broken-back shrimp), Lysmatidae (cleaner and peppermint shrimp), Oplophoridae (bioluminescent shrimp), Palaemonidae (freshwater and symbiotic anemone shrimp), and Thoridae (anemone shrimp), their last common ancestor and all of its descendants. Based on previous molecular trees, and our topology, Procarididea (anchialine shrimp) are excluded from the otherwise monophyletic crown Caridea [65]. Although we did not sequence Amphionidacea, recent analysis of four genes [66] placed this organism as a suspected larval pseudo-taxon within our concept of Caridea, thus the calibration node would not need to be modified if they are added in future. ***Fossil specimens.** Blaculla haugi* Winkler 2015 [67], holotype SMNS (Staatliches Museum für Naturkunde, Stuttgart, Germany) 70286. ***Phylogenetic justification.** B. haugi* is within crown Caridea based on overlap of the pleonal pleurae, and the first two pereiopods being chelate while the third, fourth, and fifth are achelate [67]. The first two pereiopods have some characteristics in common with the extant clade ‘Alpheoidea’ (see below), with stouter first pereiopods and multi-jointed second pereiopods [67]. However, pereiopod appendage morphology across caridean families appears to be convergent [68], which challenges the ability to correctly assign fossils to crown families based on these morphologies (also challenging the confidence in older putative caridean fossils). Mandibular structure is possibly more diagnostic [68], however, mouthparts are not well preserved in Solnhofen material [67]. Furthermore, the assumed composition of crown Alpheoidea does not include Palaemonidae [69,70], but these are found within a paraphyletic group within our tree (also supported in previous molecular trees: [71,72]). Additional putative caridean fossils are also known from the Solnhofen limestones, e.g. [73–75], and are thus of the same age, suggesting Caridea had begun to diversify. We remain agnostic on proposed, but not diagnostic, caridean fossils from the Early Triassic Paris Biota of Idaho [76]. The several possible Solnhofen taxa, together with the morphology of *B. haugi*, permit a conservative minimum age constraint on crown Caridea in its entirety, rather than on any specific families. ***Age justification.*** The fossil of *B. haugi* was found in the Solnhofen Plattenkalk (lithographic limestone) of Eichstätt, Bavaria, Germany [67]. As discussed by Benton et al. ([77], node 31), a minimum age for Solnhofen fossils is 150.94 Ma. Soft maximum as in node 2 herein.
6. ***Node.*** This node represents crown Reptantia. In our phylogeny, this is the clade comprising Axiidea, Astacidea, Achelata, Polychelida, Gebiidea, Anomura, and Brachyura, their last common ancestor and all of its descendants. Glypheidea are included in previous formulations of Reptantia, and if corroborated by future phylogenomic data (as already suggested in the mitogenome tree of Tan et al. [78]), they will not affect our calibration choice. As in the discussion of Wolfe et al. [44] node 49, the Devonian fossil *Palaeopalaemon newberryi* Whitfield 1880 [79] is within crown Reptantia. This is implicitly confirmed in the phylogenetic analysis of Jones et al. [80]. Thus all calibration data including both age priors as in Wolfe et al. [44], nodes 49, 55, and 56.
7. ***Node.*** This node represents crown Achelata, a clade comprising the families Palinuridae (spiny lobsters) and Scyllaridae (slipper lobsters), their last common ancestor and all of its descendants. ***Fossil specimens.** Yunnanopalinura schrami* Feldmann et al. 2012 [81]. Holotype LPI (Luoping section, Invertebrate Paleontology Collection, Chengdu Institute of Geology and Mineral Resources) 40169. ***Phylogenetic justification.*** This species has been previously justified as a calibration for Achelata [81,82]. ***Age justification.** Y. schrami* was recovered from limestone of the Luoping Biota, Member II of the Guanling Formation, near Luoping, Yunnan, south China [81,83]. Extensive stratigraphy of the Luoping biota places the Guanling Formation just below the uppermost boundary of the Anisian stage [83]. The upper boundary of the Anisian is estimated at 241.5 Ma ± 1 Myr [58], thus providing a minimum age at 240.5 Ma. A soft maximum age is obtained by phylogenetic bracketing, with the generous assumption that crown Achelata are not older than the oldest known crown Decapoda, which is *Palaeopalaemon newberryi* [44]. The age of the Chagrin Shale, which bears *P. newberryi*, is late Fammenian based on the presence of index algae; there is no lower bound index fossil mentioned [44]. Thus a maximum age of this deposit is estimated as the lower bound of the Fammenian, at 372.2 Ma [84].
8. ***Node.*** This node represents crown Nephropidae (true lobsters). In our tree, this includes *Thaumastocheles*, *Nephropoides*, and *Nephropsis*, their last common ancestor and all of its descendants. Note that the family Thaumastochelidae is now synonymized with Nephropidae [85]. ***Fossil specimens.** Jagtia kunradensis* Tshudy & Sorhannus 2000 [86]. Holotype IRScNB (Institut Royal des Sciences Naturelles de Belgique, Brussels) 90-33h. ***Phylogenetic justification.*** Morphological phylogenetic analysis places *J. kunradensis* as more closely related to *Nephropsis* and *Nephropoides* than to *Thaumastocheles* [87]. Thus *J. kunradensis* is in the crown group. Another lobster fossil (*Oncopareia*) within the crown group in this analysis was not referred to specific material, and is of similar age to *J. kunradensis. Hoploparia stokesi* was also found within the crown group Nephropidae by Karasawa et al. [87], but the systematics of this genus demand revision [88]. Such revisions would likely compromise the wider stratigraphic range reported for *Hoploparia* [89]. Thus *J. kunradensis* is the most conservative calibration fossil for Nephropidae. ***Age justification.** J. kunradensis* has been collected from the Kunrade Limestone facies of the Maastricht Formation, southeast Netherlands [86]. The Maastricht Formation is eponymous for the Maastrichtian stage of the latest Cretaceous (although it does not bear the GSSP for either lower or upper stage boundary; [64]). The upper boundary of the Maastrichtian is well constrained at 66.0 Ma [90], thus providing a minimum age for Nephropidae. Soft maximum as in node 8 herein.
9. ***Node.*** This clade, in our tree, comprises Cambaridae, Cambaroididae, Astacidae, and Parastacidae (together: crayfish), their last common ancestor and all of its descendants. Monophyly of freshwater crayfishes has been previously established by a number of molecular and morphological analyses, e.g. [82,91]. ***Fossil specimens.** Cricoidoscelosus aethus* Taylor et al. 1999 [92]. Based on holotype NIGP (Nanjing Institute of Geology and Palaeontology) 126337 and NIGP 126355 [87]. ***Phylogenetic justification.*** Morphological phylogenetic analysis places *C. aethus* as the sister of a clade comprising the extant crayfish *Cambarus* and *Procambarus*, both members of Cambaridae [87]. The published morphological tree places *C. aethus* further crownward than Parastacidae [87], in relationships that mirror our AHE topology. Therefore, it is an appropriate fossil to calibrate the freshwater crayfish crown group. ***Age justification.*** Minimum as in Wolfe et al. [44], node 60. Soft maximum as in node 8 herein.
10. ***Node.*** This node represents crown Axiidea (mud shrimp/ghost shrimp). In our tree, this clade is represented by Axiidae, Callianassidae, Callianideidae, Gourretiidae, their last common ancestor and all of its descendants. Monophyly of this clade is established by molecular phylogeny of the 16S, 28S, and 18S genes [93,94]. ***Fossil specimens.** Protaxius isochela* Woodward 1876 [95]. Calibration material is from specimens MAN (Museum-Aquarium at Nancy, France) 11700-11762 [96]. ***Phylogenetic justification.*** As discussed by Hyžný & Klompmaker [97], the fossil record of mud shrimps usually preserves only the distal cheliped elements, and many are assumed to be members of a wastebasket ‘*Callianassa*’, which compromises systematic identification. The oldest putative members are Jurassic, all of which are likely most closely related to crown Axiidae [97]. The very oldest, *Magila bonjouri* Étallon 1861, was described from a preserved dactylus + propodus, but the holotype cannot be found [97]. The only Jurassic species with full body preservation is *P. isochela*, where the specimen figured (drawn) by Woodward is unfortunately not identified ([97], supplement). However, Hyžný & Klompmaker [97] mention material of *P. isochela* from France as accepted within this taxon, thus we may calibrate a minimum age from the specimens discussed by Breton et al. [96]. Although the relationship of *P. isochela* to crown Axiidae is not precisely known, it is well within the crown-group of Axiidea. ***Age justification.*** French *P. isochela* material was recovered from wells at Bure, Meuse, in the northeast of France [96]. The locality belongs to the *Rasenia cymodoce* to *Aulacostephanus mutabilis* ammonite Zone [96]. The upper boundary of the *A. mutabilis* Zone is Chron M23r.2r.1, with an age of 153.55 Ma [98], in the Kimmeridgian. This provides a minimum age estimate for Axiidea. Soft maximum as in node 8 herein.
11. ***Node.*** This node represents crown Gebiidea (mud shrimp/mud lobster/ghost shrimp). In our tree, this is only represented by Laomediidae and Axianassidae, their last common ancestor and all of its descendants. ***Fossil specimens.** Laurentiella imaizumii* Karasawa 1993 [99]. Material figured by Karasawa [99] includes MFM (Mizunami Fossil Museum, Mizunami, Gifu Prefecture, Japan) 39003-39006, and MFM 39117. ***Phylogenetic justification.*** Based on molecular phylogenetics, Laomediidae and Axianassidae are sister clades within Gebiidea, exclusive of the clades we did not sample: Thalassinidae and Upogebiidae [93,94]. No fossil Axianassidae are recorded, thus we must calibrate based on fossil Laomediidae. While the position of *L. imaizumii* within either the crown or stem of Laomediidae is unknown, it does share characters indicating its membership within the total group of Laomediidae (e.g. strongly heterochelate chelipeds; [99]). Thus, *L. imaizumii* is within the crown group of the represented Gebiidea. Note that the extant genus *Laurentiella* is considered a junior homonym of *Saintlaurentiella* [100]; this does not influence the calibration choice. ***Age justification.*** The oldest occurrence of *L. imaizumii* is in the Akeyo Formation of the Mizunami Group, Gifu Prefecture, Japan [99,101]. The type locality, the Toyoda Formation, is slightly younger [99,102]. Thus we calibrate based on the Akeyo Formation, which is the uppermost member of the Mizunami Group, and correlated to the C5Dr chron based on diatom fossils [101]. Globally, these strata underlie the correlated NMU 5 (Asia) and MN 5 (Europe) units. Thus a conservative upper bound age for the Akeyo Formation is 17 Ma. Soft maximum as in node 8 herein.
12. ***Node.*** This clade, in our tree, comprises Porcellanidae (porcelain crabs) and Munididae (some squat lobsters), their last common ancestor and all of its descendants. Based on previous total evidence analysis [103], these families are close relatives. The full breadth of Galatheoidea are not represented here. ***Fossil specimens.** Jurellana tithonia* Schweitzer & Feldmann 2010 [104]. Holotype NHMW (Natural History Museum of Vienna) 1990/0041/2518 and paratype NHMW 1990/0041/1445. ***Phylogenetic justification.** J. tithonia* is known from dorsal carapace material, and was previously used to calibrate Porcellanidae [103]. Its carapace is flattened and carcinized, with a well developed cervical groove, and overall poorly defined regions [104]. The carapace shape excludes *J. tithonia* from the crown group of any other galatheoid clades. It may be distinguished from brachyurans based on the carapace flank being relatively short [104]. These characters together confirm that *J. tithonia* is at the least, a member of the total group of Porcellanidae, and thus belongs within the crown group of Porcellanidae + Munididae. Putative members of Munididae [105] and Munidopsidae [106] were also discovered in the Ernstbrunn Limestone. These discoveries further support divergence of the major Galatheoidea lineages prior to the Cretaceous. ***Age justification.** J. tithonia* was recovered from the Ernstbrunn Limestone, lower Austria [104]. Ammonites are unavailable for precise biostratigraphy from this locality, but a nearby locality from the same unit preserves the ammonites *Richterella richteri*, *Simplisphinctes*, and the diagnostic *Micracanthoceras microcanthum* constraint for the late Tithonian [98,107]. However, benthic foraminifera and calcareous algae suggest the Ernstbrunn Limestone may have been deposited as late as the early Cretaceous, early Berriasian stage [108]. Although most literature accepts a Tithonian age, we recognize that the more conservative constraint should allow some probability density in the early Berriasian, which has a minimum age of approximately 142 Ma [98]. A soft maximum age is obtained by phylogenetic bracketing, with the generous assumption that crown Porcellanidae + Munididae are not older than the oldest putative crown Anomura. Previous studies have suggested the oldest Anomura is *Platykotta akaina* [103,109]. However, its second pair of chelate pereiopods makes this affinity uncertain and it could even be stem-group Meiura [110]; nevertheless, *P. akaina* is surely older than crown Porcellanidae + Munididae. *P. akaina* was collected at Wadi Naqab close to Ras Al Khaimah City, United Arab Emirates, from limestone of the Ghalilah Formation [109]. No precise constraint is available, so a soft maximum age is the base of the Norian stage, at ~227 Ma.
13. ***Node.*** This node represents crown Paguroidea (hermit and king crabs). In our tree, this includes Coenobitidae, Diogenidae, Paguridae, Lithodidae, their last common ancestor and all of its descendants. Our AHE results follow the topology of the Bayesian molecular-only analysis of Bracken-Grissom et al. ([103], their Figure 2) in excluding Parapaguridae from a monophyletic Paguroidea. ***Fossil specimens.** Diogenicheles theodorae* Fraaije et al. 2012 [111]. Holotype I–F/MP/3957/1533/08 (Institute of Systematics and Evolution of Animals, Polish Academy of Sciences, Kraków, Poland). ***Phylogenetic justification.*** The assignment of *D. theodorae* to crown Paguroidea is based on carapace material, and was justified in Bracken-Grissom et al. [103]. It bears similarities (a distinct threefold junction of keraial, massetic and anterior branchial areas of the outer carapace) with the family Parapylochelidae, which is likely closely related to the other symmetrical hermit crabs in Pylochelidae [111]. The total evidence phylogeny of Bracken-Grissom et al. [103] places Pylochelidae as the most deeply branching lineage within Paguroidea. ***Age justification.** D. theodorae* was discovered in an abandoned quarry in Bzów, southern Poland [111]. As discussed by Fraaije et al. [111], this locality preserves the ammonites *Ochetoceras canaliculatum*, *Trimarginites trimarginatus*, *Dichotomosphinctes* sp., and *Glochioceras subclausum.* Together, these ammonites are globally correlated to the *Gregoriceras transversarium* ammonite Zone of the Oxfordian. Cyclo- and magnetostratigraphy indicate a minimum age of 159.44 Ma [98] for the *G. transversarium* Zone, and thus for *D. theodorae.* Soft maximum as in node 13 herein.
14. ***Node.*** This clade, in our tree, comprises Paguridae (hermit crabs) and Lithodidae (king crabs), their last common ancestor and all of its descendants. ***Fossil specimens.** Paralithodes bishuensis* Karasawa et al. 2017 [112]. Holotype MFM 83077. ***Phylogenetic justification.*** No phylogenetic analysis has yet been conducted to evaluate the relationships among extant and fossil pagurids, and the taxonomy of fossil members remains problematic [113]. As Paguridae is paraphyletic with respect to Lithodidae in previous molecular and total evidence analyses [103,114], fossils that may be within Paguridae are not guaranteed to fall within this node (as they may be members of lineages leading to pagurids that are outside of our AHE taxon sampling). Therefore we conservatively use the oldest likely fossil of Lithodidae. *P. bishuensis* possesses several diagnostic features allying it with members of the extant genus *Paralithodes*, particularly the sparsely arranged low, pointed dorsal tubercles on the carapace [112]. Thus, a position within crown Lithodidae is confirmed. Although previous publications have suggested the oldest Lithodidae is *Paralomis debodeorum* Feldmann 1998 [115], the sediments in which it is found are poorly constrained and may be as young as the Pliocene. ***Age justification.*** The type locality for *P. bishuensis* is ‘locality MRZ06’, Minamichitacho, Aichi Prefecture, Japan, found in sandstone of the Yamami Formation of the Morozaki Group [112]. Biostratigraphic index fossils include the diatom *Crucidenticula sawamurae*, which is correlated in North America to the C5Cn chron, ECDZ2, and *Delphineis ovata* Zone [116]. The approximate age of the top of the ECDZ2 is 15.8 Ma [116], providing a minimum age constraint. Soft maximum as in node 13 herein.
15. ***Node.*** This node represents crown Eubrachyura (‘higher’ true crabs). In our tree, this includes Heterotremata and Thoracotremata, their last common ancestor and all of its descendants. Monophyly of Eubrachyura has been supported in previous molecular analyses [117]. ***Fossil specimens.** Telamonocarcinus antiquus* Luque 2015 [118]. Holotype IGM (Colombian Geological Survey, Bogotá, Colombia) p881012. ***Phylogenetic justification.*** The phylogenetic position of *T. antiquus* within Eubrachyura is based on characters shared with extant Dorippoidea (i.e. Dorippidae and Ethusidae). While the position of gonopores is unclear [118], preventing definitive assignment to Heterotremata, the distinctive carapace outline and groove pattern in particular ally this fossil with crown Dorippoidea. Luque [118] concluded that Dorippoidea may be an early branching lineage of Eubrachyura. Although no members of Dorippidae or Ethusidae were included in our AHE sampling, their closest relative in a previous molecular phylogeny was Leucosiidae [117], which we did include. Given the uncertainty in the fossil’s possession of the defining character of Heterotremata (male coxal gonopores), and the exact position of Dorippoidea, we conservatively calibrate only the Eubrachyura using *T. antiquus*. *Age justification.* The fossil of *T. antiquus* was discovered from shales of the lowermost Tablazo Formation, in El Batán, Montegrande, near the town of La Fuente, Department of Santander, Colombia [118]. The Tablazo Formation bears *Parahoplites* and *Douvilleiceras* ammonites, and locally is within the *Douvilleiceras solitae*–*Neodeshayesites columbianus* Zone [118]. Globally, *Douvilleiceras mammillatum* straddles the early to middle Albian [64]. The *D. mammillatum* Zone thus provides a conservative minimum age for *T. antiquus* at 110.22 Ma [64]. A soft maximum age is obtained by phylogenetic bracketing, with the generous assumption that crown Eubrachyura are not older than the oldest crown Brachyura. The oldest crown Brachyura is debatable; *Eocarcinus praecursor* [119] and *Eoprosopon klugi* [120] have both been proposed, but both lack some crown characters and are only represented by rather poorly preserved dorsal carapaces [110]. Nevertheless, stem-lineage positions of these taxa would imply the Brachyura crown may be even younger, so we calibrate the soft maximum from the base of the Pliensbachian, at 191.8 Ma.
16. ***Node.*** This node represents crown Thoracotremata. In our tree, this clade is comprised of the sampled families Grapsidae (marsh/shore crabs), Ocypodidae (ghost and fiddler crabs), Plagusiidae, Sesarmidae, and Varunidae, their last common ancestor and all of its descendants. Monophyly of Thoracotremata is supported by male gonopores located on the sternum, and by previous molecular phylogenies [117]. Monophyly of previously discussed superfamilies (e.g. Grapsoidea, Ocypodidea) is under suspicion from this and other molecular phylogenies [117,121], so the clade treated herein remains Thoracotremata. ***Fossil specimens.** Litograpsus parvus* Müller & Collins 1991 [122] (as revised by Schweitzer & Karasawa 2004 [123]), holotype M.91-227 (Natural History Museum of Hungary). ***Phylogenetic justification.*** Members of Thoracotremata are rare and hard to identify in the fossil record, likely because many live in difficult to preserve intertidal and semiterrestrial habitats. *L. parvus* shares characters with extant Grapsidae and Sesarmidae, such as the rectangular carapace, size and positioning of the orbits, and a transverse ridge formed by the cardiac region with broad branchial ridges [123]. Although the exact relationship to the extant members is unknown, based on our topology, a position of *L. parvus* on the stem of either Grapsidae or Sesarmidae would still be within crown-group Thoracotremata. Although without sufficient confirmation, some older crown-group fossils most likely exist for Thoracotremata. There is a possible ‘grapsoid’ crab from mid-late Paleocene sediments of Colombia (Luque et al. 2017 [124], Fig. 8J), but it has not yet received systematic study. Possible stem-group ‘Pinnotheridae’ fossils *Viapinnixa alvarezi* and *V. perrilliatae* are known from the early Eocene (Ypresian; [125,126]) of Chiapas, Mexico; however Pinnotheridae are not sampled here, and at least some members may fall outside of the crown-group we define [117,127–129]. Finally, limb fragments of *Varuna*? sp. have been reported from the middle Eocene (Lutetian?) of Jamaica [124,130,131], but with limited specimen or stratigraphic information. ***Age justification.** L. parvus* is known from limestone sediments of the Szépvölgy Formation, Hungary [122]. Co-occurring foraminifera constrain the age of the Szépvölgy Limestone to the NP20 zone of the C15n chron [122,132]. This is Priabonian, with an upper boundary of 33.9 Ma, providing a minimum age constraint. Soft maximum as in node 16 herein.
17. ***Node.*** This node represents crown Heterotremata. In our tree, this clade is comprised of the sampled families Atelecyclidae, Bellidae, Corystidae, Leucosiidae (purse crabs), Menippidae (stone crabs), Platyxanthidae, and the superfamilies Portunoidea (swimming crabs), Xanthoidea (mud crabs) and Majoidea (spider and decorator crabs), their last common ancestor and all of its descendants. Composition of Portunoidea as in Evans [133]. Compositions of Xanthoidea and Majoidea are as defined in the molecular analysis of Tsang et al. [117], although monophyly of their constituent families remains questionable. ***Fossil specimens.** Cretamaja granulata* Klompmaker 2013 [134]. Holotype MGSB (Museo Geológico del Seminario de Barcelona, Spain) 77706A+B. ***Phylogenetic justification.*** Klompmaker [134] diagnosed *Cretamaja* as appropriately belonging to Majoidea based on carapace shape (which exhibits rampant convergence among brachyurans) and presence of anterolateral spines. These characters (especially because of the limitations of carapace shape) only permit assignment to a deeply divergent lineage of Majoidea, however, likely outside the Majoidea crown group [117]. While monophyly of Majoidea has been supported by previous molecular and morphological phylogenies [117,135–137], the exact relationships among constituent families (in our tree, Epialtidae, Inachoididae, and Mithracidae) are debated. A position for *C. granulata* along the stem of Majoidea would still permit assignment to the crown group of Heterotremata, and thus a calibration of the latter clade. ***Age justification.*** The Koskobilo fauna belongs to the Albinez Unit of the Eguino Formation, southwest of Alsasua, Spain [134]. Either a Cenomanian or late Albian age has been discussed for the Ablinez Unit, based on underlying ammonites and those of contemporaneous reef deposits (summarized by Klompmaker [134]). *Mortoniceras perinflatum* was one of the contemporaneous ammonites from a nearby locality, and it is an index fossil for the late Albian [64], a convincing age. The upper boundary of the *M. perinflatum* Zone is at 100.91 Ma [64], which is therefore the minimum age of *C. granulata.* Soft maximum as in node 16 herein.
18. ***Node.*** This node represents crown Majoidea (spider and decorator crabs). In our tree, this clade includes Epialtidae, Inachoididae, and Mithracidae, their last common ancestor and all of its descendants. While monophyly of Majoidea has been supported by previous molecular and morphological phylogenies [117,135–137], the exact relationships among constituent families are debated. ***Fossil specimens.** Planobranchia palmuelleri* Artal et al. 2014 [138]. Holotype MGSB 79782. ***Phylogenetic justification.*** The most distinctive characters for majoids are the carapace shape. *P. palmuelleri* has an advanced carapace front with straight lateral margins, rounded longitudinal frontal ridges, lateral orbits with a strong outer-orbital subtriangular tooth, and dorsal conical spines: these characters refer *Planobranchia* to the Inachidae. Membership on the stem-lineage of Inachidae tentatively confirms the ability to provide a minimum calibration for the crown Majoidea we have sampled. Late Cretaceous [124,139], Eocene [140,141], and Oligocene [142] putative majoids have been discovered with preserved carapaces, but they either fall outside of our molecular crown taxon sampling, or lack specimen information. ***Age justification.** P. palmuelleri* is known from strata of the Vic area, Barcelona province, Catalonia, Spain, most likely assigned to the Coll de Malla Formation [138]. As there is some controversy over the precise lithostratigraphic unit [143], we agree with Artal et al. [138] that a Lutetian age is conservatively appropriate. The upper bound of the Lutetian is 41.2 Ma, providing a minimum age. Soft maximum as in node 16 herein.
19. ***Node.*** This node represents crown Xanthoidea (mud crabs). In our tree, the members are ‘Xanthidae’ and Panopeidae, their last common ancestor and all of its descendants. Monophyly of a clade containing at least these members of Xanthoidea is supported by previous molecular and total evidence phylogenies [117,144]. ***Fossil specimens.** Phlyctenodes tuberculosus* Milne Edwards 1862 [145]. The holotype MNHN (Muséum National d’Histoire Naturelle, Paris) R03826, and specimen MCZ (Museum of Comparative Zoology, Harvard University) 2456 are figured by Busulini et al. [146]. ***Phylogenetic justification.** Phlyctenodes* fossils are only known from carapaces, which are linked with members of the xanthid subfamily Actaeinae based on the ornamentation of the carapace with tubercles [146,147]. However, as many of the subfamilies within Xanthidae, including Actaeinae, are very likely polyphyletic [144,148], and even our focal analysis suggests that Xanthidae may be paraphyletic, the exact relationship of *P. tuberculosus* to sampled taxa is unclear. Thus we calibrate the crown group of all of sampled Xanthoidea. Of contemporaneous Xanthoidea fossils [147], *P. tuberculosus* has recently refigured and discussed specimens, and is thus selected. ***Age justification.*** The holotype of P. *tuberculosus* was attributed to the locality Hastingues, Landes, France, in the ‘middle Eocene’ [146]. The MCZ specimen was discovered in the better known San Feliciano Hill quarry of the Berici Hills, Vicenza, Italy [146]. The decapod-bearing strata of the quarry are correlated to the lower Priabonian stage, late Eocene, based on calcareous nannofossils [149,150]. The upper boundary of the Priabonian is 33.9 Ma, providing a minimum age constraint. Soft maximum as in node 16 herein.

---

**Figure S1.**
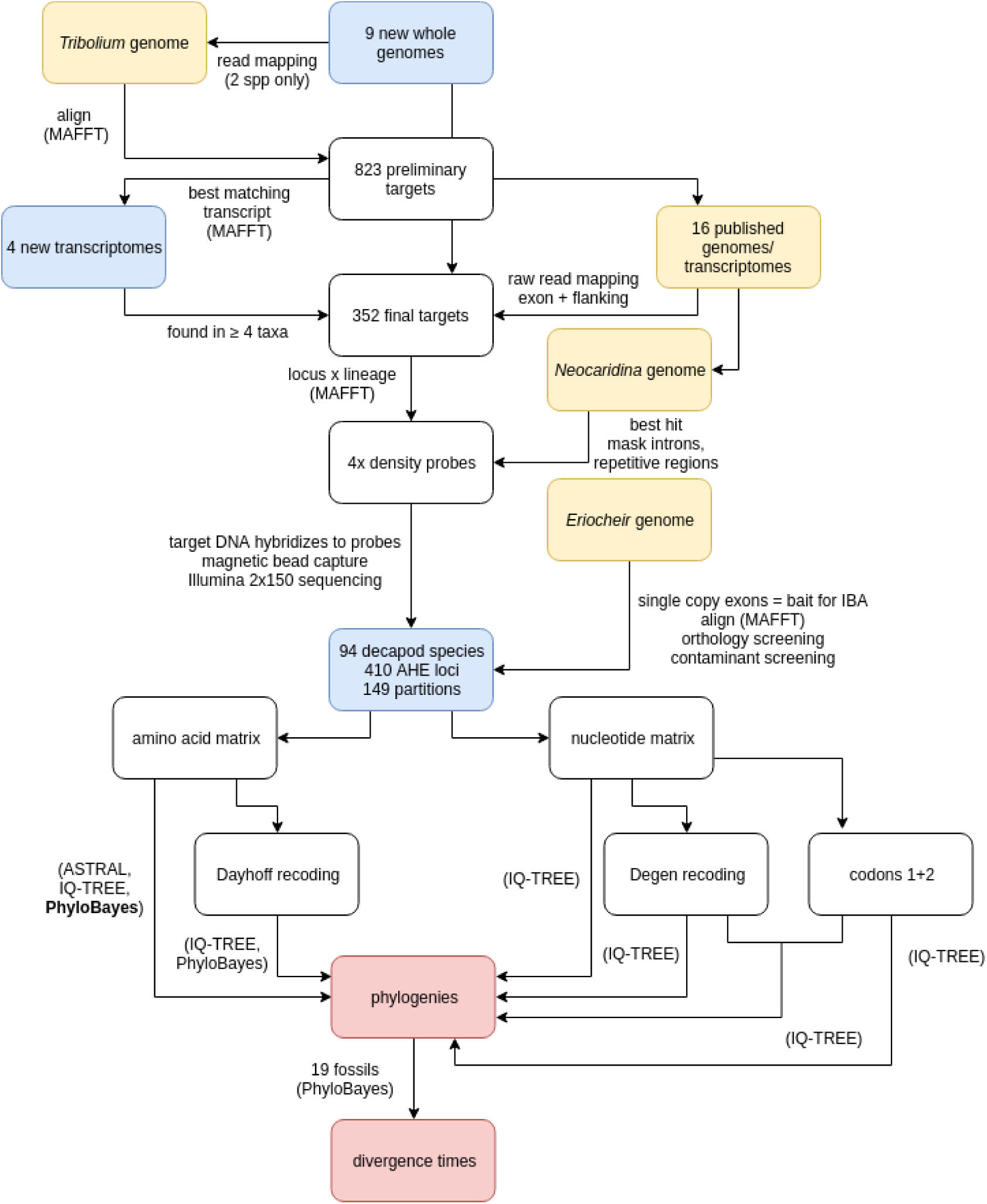
Workflow for our AHE data collection and analysis. Yellow boxes represent published genomic resources, blue boxes are newly sequenced in this paper, and red boxes are the focal results in **Figures 2-3**.

**Figure S2.**
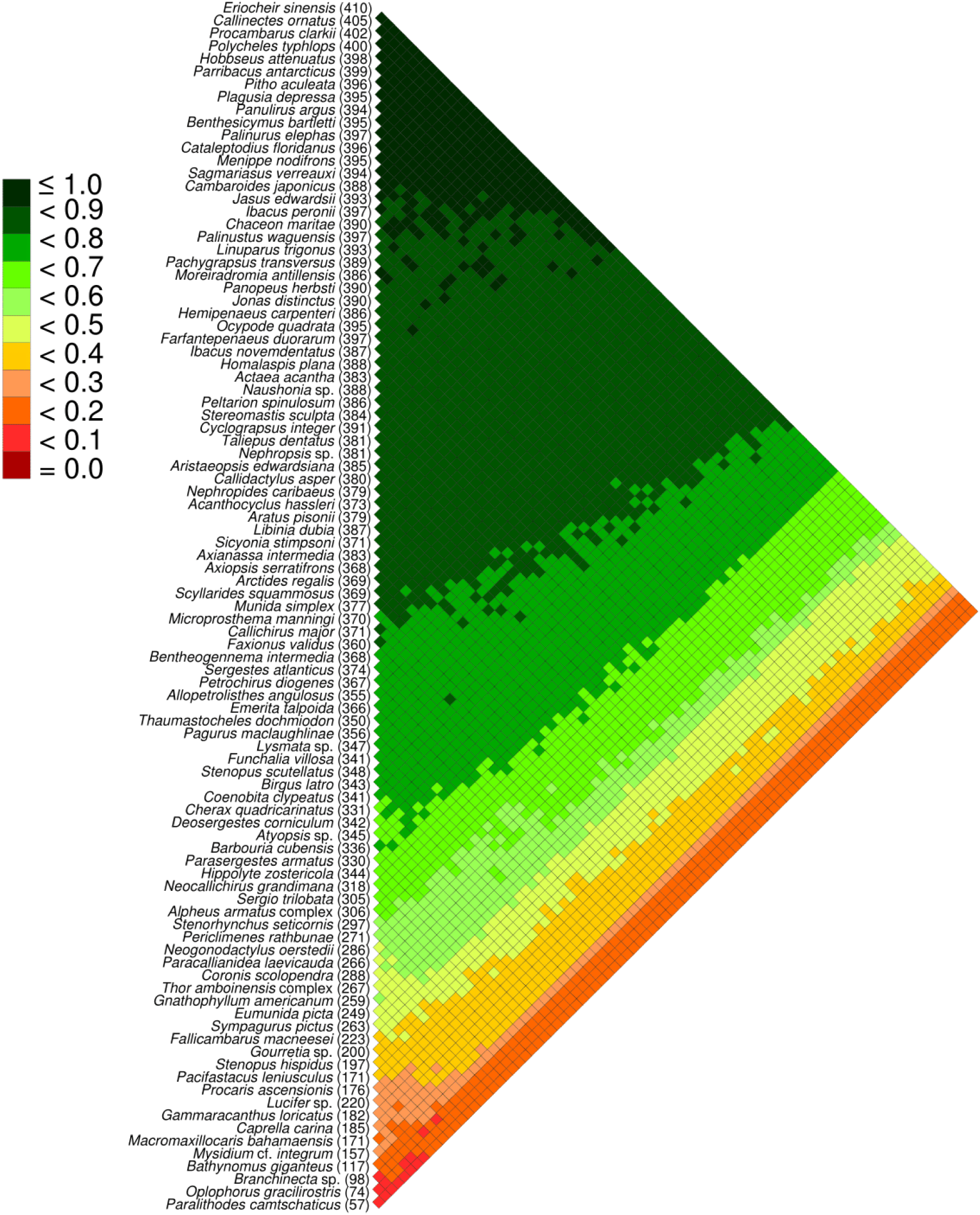
Pairwise heat map of species-pairwise amino acid dataset completeness for all targeted AHE loci, in the unrecoded amino acid dataset. Numbers in parentheses are total captured loci per species. Low shared site coverage in shades of red and high shared site coverage in shades of green.

**Figure S3.**
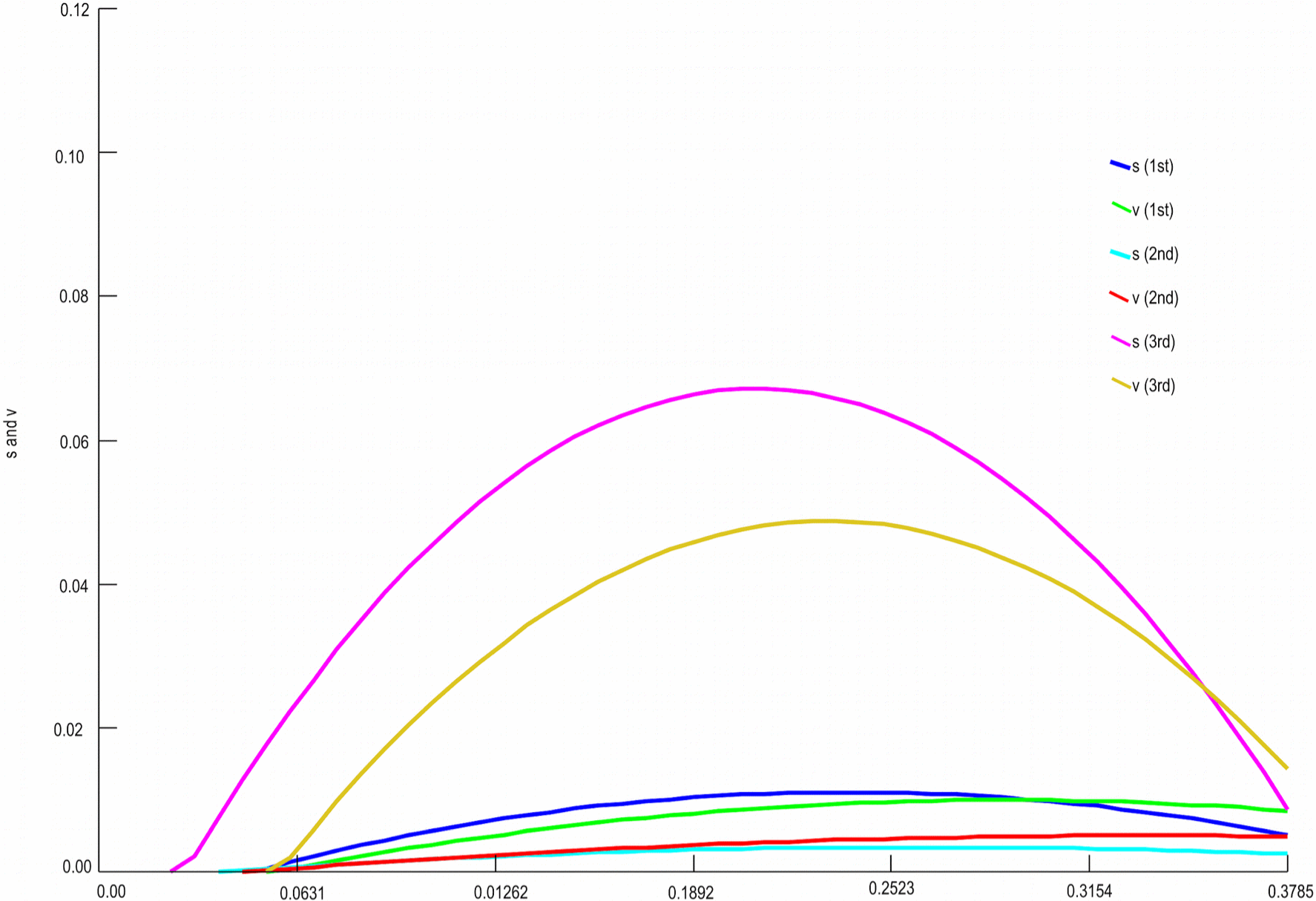
Saturation plot for each codon position with transversions (v) and transitions (s) plotted against F84 distance. The third codon position clearly deviates from expected values, and thus has experienced saturation.

**Figure S4.**
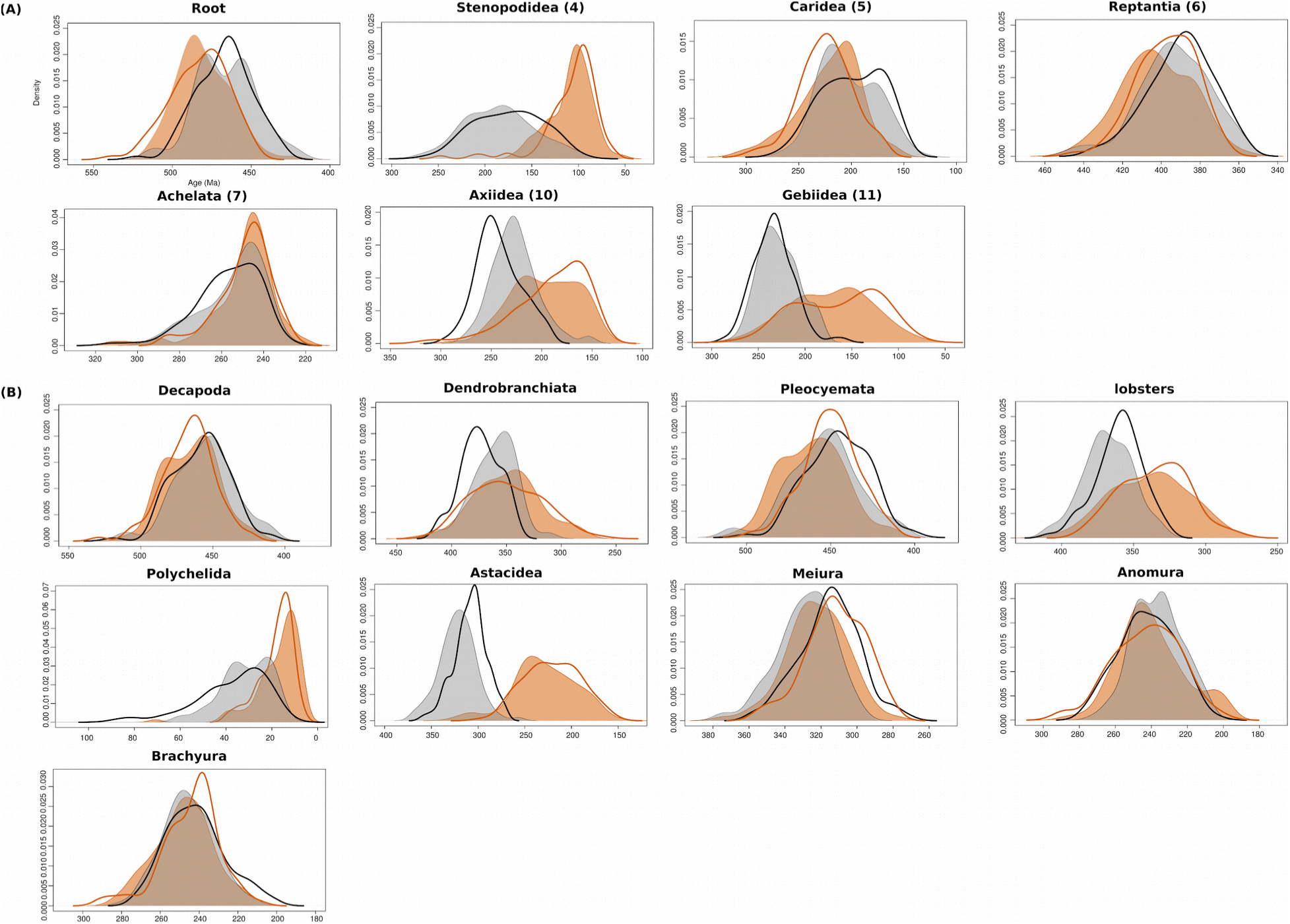
Comparison of posterior probability distributions for divergence times assessed as in Figure 3 (posterior), and using the same analyses under the effective prior (removing sequence data). The posterior analyses are shaded; effective priors are superimposed on the same axes with a heavy line of the same color. Grey/black analyses with the CIR autocorrelated clock model (depicted in Figure 3); orange analyses with the UGAM uncorrelated clock model. *(a)* Selected nodes directly calibrated by fossils and their calibration number; *(b)* Selected nodes calibrated by only a birth-death tree prior.

All Extended Tables are available as.xslx or.csv files attached.

**Table S1.** Details of all transcriptome and genome sequences used in probe design.

**Table S2.** Sample information for whole genome sequencing.

**Table S3.** Sample information for transcriptome sequencing.

**Table S4.** Assembly statistics for transcriptome sequencing.

**Table S5.** Brief description of enrichment kits for each of six selected major lineages (Achelata, Anomura, Astacidea, Brachyura, Caridea, and Dendrobranchiata).

**Table S6.** Sample information for AHE sequencing.

**Table S7.** Formatted list of node calibration priors.

**Table S8.** Assembly statistics for AHE sequencing, and loci sequenced for each species. For each locus, 1 represents presence and 0 represents absence in the main data matrix.

